# Palantir characterizes cell fate continuities in human hematopoiesis

**DOI:** 10.1101/385328

**Authors:** Manu Setty, Vaidotas Kiseliovas, Jacob Levine, Adam Gayoso, Linas Mazutis, Dana Pe’er

## Abstract

Recent studies using single cell RNA-seq (scRNA-seq) data derived from differentiating systems have raised fundamental questions regarding the discrete vs continuous nature of both differentiation and cell fate. Here we present Palantir, an algorithm that models trajectories of differentiating cells, which treats cell-fate as a probabilistic process, and leverages entropy to measure the changing nature of cell plasticity along the differentiation trajectory. Palantir generates a high resolution pseudotime ordering of cells, and assigns each cell state with its probability to differentiate into each terminal state. We apply Palantir to human bone marrow scRNA-seq data and detect key landmarks of hematopoietic differentiation. Palantir’s resolution enables identification of key transcription factors driving lineage fate choices, as these TFs closely track when cells lose plasticity. We demonstrate that Palantir is generalizable to diverse tissue types and well-suited to resolve less studied differentiating systems.

## Introduction

Differentiation is among the most fundamental processes in biology. The traditional view of cell differentiation comprises a series of discrete steps through well-defined stages, through which a cell transitions from less-differentiated to a more differentiated state. Single cell studies ^1–6^ have however demonstrated that in differentiating systems, cell states are observed to reside along largely continuous spaces. Despite the evolution in thinking about differentiation as a continuous process, cell fate choices continue to be largely viewed as a series of discrete bifurcations, along a developmental trajectory, towards terminal cell states ^7–9^.

Contrary to this view, studies based upon epigenomic measurements such as DNase-seq and ATAC-seq indicate that progressive restriction of the enhancer landscape coupled with pre-establishment of lineage specifying enhancers in precursors can serve as vehicle for driving differentiation, suggesting mechanisms that might underlie a continuous process ^5, 10–13^. Indeed, we observe a lack of well-defined bifurcation points when scRNA-seq profiles are projected along the strongest axes of variation (Fig. 1a). Even at the level of individual genes, we observe a broad representation of observed gene ratios, rather than bimodal expression states (Fig 1a). In support of cell fate choice continuity, a recent study profiling trans-differentiation in mice demonstrated that individual clones do not deterministically reach particular differentiated states, but rather fate choices are inherently probabilistic in nature ^14–16^. These data raise fundamental questions concerning cell fate determination. Are cell fates, like cell state transitions, continuous? When and how might cell fate choices be made in a continuous model?

**Fig 1:**
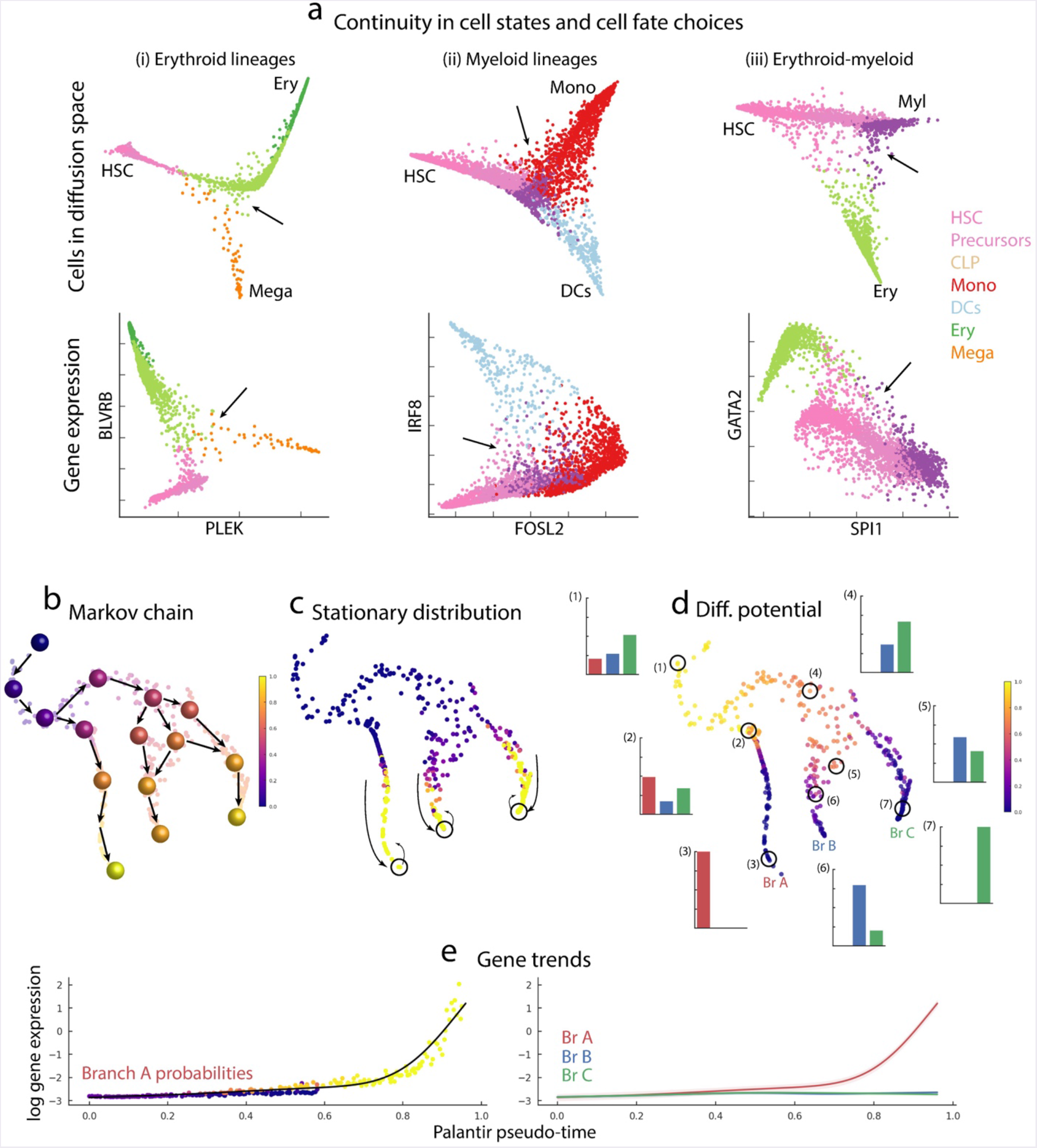
Palantir determines continuities in cell state transitions and cell fate choices in differentiation. (a) Top panel: Plots showing the projection of CD34+ cells from human bone marrow along diffusion components. Cells are colored by Phenograph clusters (Supp. Fig. 4a). Arrows highlight the decision making regions and show the pervasive lack of well-defined branch points, highlighting the continuity in cell fate choices. Bottom panel: Scatter plots showing gene expression for gene pairs involved in lineage decisions for the same cells in the corresponding top panel. (b-d) A cartoon illustration of Palantir constructs and outputs, based on toy subsampled dataset from CD34+ human hematopoiesis data. Plotted is the phenotypic manifold: each dot represents a cell, embedded into diffusion space, based on the first 3 components and visualized using tSNE applied to the embedded space. (b) Cells are colored based on pseudo-time, with a cartoon illustration of constructing a Markov chain over the phenotypic manifold. (c) Each cell is colored based on the stationary distribution of the Markov chain constructed in (b) demonstrating clear outliers in the mature states. Outliers of the stationary distribution that are also boundary states (circles) are selected as terminal states. (d) Cells colored based on differentiation potential. Specific examples are highlighted to show the relationship between pseudo-time, differentiation potential and branch probabilities: For each highlighted cell, a histogram represent the branch probabilities, color coded in a manner corresponding to each of the 3 terminal states. Earliest cells have the highest differentiation potential (1) with a gradual decrease as cells move towards commitment (2-3). Modeling of cell fate choices as probabilities provides a representation of the continuity in cell fate choices (4-7). (e) Plot showing the expression of a branch A specific gene along pseudo-time. Each dot represents a cell, with the x-axis representing pseudo-time, y-axis the gene expression and colored by probability of the cell reaching branch A. The black line represents the computed gene expression trend for this data (left panel). (Right panel) The expression trend for the same gene plotted for each of the 3 lineages. The unified framework of pseudo-time and branch probabilities enable characterization of gene expression dynamics across a common axis.

Our approach to investigating these fundamental questions is to leverage single cell RNA-seq (scRNA-seq) data to model the landscape of differentiation and characterize cell state and fate continuities. As differentiation is *asynchronous*, scRNA-seq derived from a population of differentiating cells yields a snapshot representing the full dynamic range of these cell states. Here we present Palantir, an algorithm that models continuity in both cell state and fate choice. Palantir takes as input scRNA-seq from a single asynchronous sample and markers representing an early cell to generate a pseudo-time ordering of cells and assigns each cell state with its probability to differentiate into each potential terminal state.

We applied Palantir to characterize human hematopoietic differentiation using scRNA-seq profiles of ~25000 cells enriched for CD34, a marker for hematopoietic stem and progenitor cells ^17, 18^. Palantir identified known terminal states of human hematopoietic differentiation and ordered cells along a pseudo-time that recapitulated known marker trends along development. Critically, Palantir identified points along the trajectory where the differentiation potential drastically shifts; these shifts mark key events in hematopoiesis. Thus, Palantir provides a quantitative approach to characterizing a continuous model of cell fate choice.

## Results

### Development as a Markov Process

Differentiation proceeds through cell divisions and progressive changes in phenotype. Because daughter cells are generally very similar to their mother cells, the population is established by incremental divergences, driven by regulatory mechanisms that create paths through the space of possible cell states. Thus, regulation constrains cell states to a low dimensional manifold of possible phenotypes ^19^. Nearest-neighbor graphs, where each node represents a particular cell state and edges connect most similar cells, have been widely used to model this manifold ^1–3, 20^.

We therefore use nearest-neighbor graphs to model differentiation in hematopoiesis. Importantly, hematopoiesis is asynchronous: a single bone marrow sample contains both the full spectrum of cell states occurring in hematopoiesis, and critically, the frequencies of each cell state. We leverage cell state frequencies to inform our model of possible and most likely differentiation paths, assuming paths in the neighbor graph represent possible differentiation paths. Critically, paths along the neighbor graph do not represent the path of a *particular* cell, but rather represent likely trajectories of cells in the population as they differentiate towards terminal states. Each cell state, represented as a node in the manifold graph, is associated with a probability distribution for reaching the terminal states. We assert that cells traverse the manifold in small steps which can be modeled using a Markov chain to represent cell fate choices in a probabilistic manner.

Two critical assumptions enable us to model differentiation as a Markov process. Firstly, differentiation progresses from a less differentiated state to a more differentiated state, i.e., cells cannot dedifferentiate into earlier states. While this assumption might be violated in some cases, we posit that it is a reasonable first order approximation, particularly in healthy differentiation. Moreover, the assumption that differentiation is unidirectional underlies all pseudo-time/trajectory detection algorithms ^1, 3, 7, 8^. We note that in aberrant systems such as cancer this assumption does not hold and other sources of information, such as mutations, are needed to guide the directionality.

A second assumption enables us to model differentiation as a Markov chain. For any node in the manifold graph, the probability of traversing to any of its neighbors is independent of its history, i.e. the path taken to reach that state. Note that for a *particular* cell, the cell’s previous developmental history is likely to be encoded in its epigenetic profile and will likely impact cell fate choices. However, nodes in the graph manifold are cell states representing multiple histories and potential trajectories rather than the developmental path of an individual cell. The distribution and frequencies of all observed cells informs the basic structure and connectivity among nodes in the graph manifold.

### The Palantir Algorithm

Given scRNA-seq collected from a sample of differentiating cells and the expression profile of an early cell, Palantir provides (1) an ordering of cells along a pseudo-time; (2) characterization of terminal differentiated states; (3) and assigns each cell a probability distribution representing the cell’s *branch probability* (BP) for reaching each of the terminal states.

First, we represent the phenotypic manifold using a nearest-neighbor graph that connects each cell to its most similar cells (Supp. Fig. 1a, Methods). Key to Palantir’s success is good graph construction so that edges connect between cells in similar developmental states and longer paths correspond to developmental trajectories. Therefore, we use diffusion maps ^21^ to focus on developmental trends and avoid spurious edges resulting from the sparsity and noise in scRNA-seq. Projecting the data onto the top diffusion components effectively focus edges in directions with high cell density and reweighs similarity along these directions (Supp. Fig. 1a).

Diffusion maps have been previously used to study differentiation in single cell data ^2, 3^ and are particularly adept at capturing differentiation trajectories ^3, 22, 23^. Typically, diffusion maps have been used to characterize pseudo-time ordering of cells by constructing separate trajectories for each major axes of variation based on individual diffusion components (DCs) ^23, 24^. While a single DC can offer a reasonable approximation of a trajectory towards a specific fate, we often observe many too many relationships between DCs and trajectories leading to each terminal fate (Supp. Fig. 2). Therefore, Palantir takes multiple DCs into account when computing the pseudo-time ordering of cells and uses shortest paths from a user defined early cell in the neighbor-graph to initiate pseudo-time. Pseudo-time is iteratively refined by means of shortest path distances from waypoints, a set of cells sampled to span the differentiation landscape (Supp. Fig. 1b-c) ^1, 2^. The computed pseudo-time does not represent a single trajectory, but rather assigns each cell their relative distance from an initial early cell, regardless of their lineage or terminal fates.

We combine the neighbor graph and pseudo-time to construct a Markov chain that models differentiation as a stochastic process, where a cell reaches one or more terminal states through a series of steps in the manifold (Fig. 1b). Pseudo-time provides directionality that is used to orient edges in the neighbor graph in a manner consistent with the ordering (Supp. Fig. 1d-e). For each directed edge we assign a transition probability, representing the probability of reaching cell *j* from cell *i* in one step. The probability of reaching a more distant cell is computed over the course of many steps and will be high if many paths connect them i.e. high probability reflects a high density of observed intermediary cell states. This is computed by exponentiating the Markov matrix *t* times, where *t* represents the length of the path (Supp. Fig. 1f). The exponentiated Markov matrix directly computes the probability of a random walk over longer paths transitioning between more distant cell states. Thus, while each single step is stochastic, at longer distances the manifold graph structure implicitly encodes the developmental trajectories.

Moreover, the Markov chain can be used to infer the terminal states directly from the data. In the Markov chain, random walks are directed towards the mature states and converge onto terminal states. Thus, Palantir identifies terminal states as boundary cells (extrema of diffusion components), that are also outliers in the stationary distribution, i.e. the states into which the random walks likely converge (Fig. 1c). Once the terminal states are identified we convert the Markov chain into an *absorbing Markov chain* by setting the terminal states as absorbing states i.e., a state with no outgoing edges.

In an absorbing Markov chain, a random walk from any state will continue until it reaches a terminal absorbing state. Each individual path through the Markov chain is associated with a probability computed as the product of the individual transition probabilities for each step taken along the path. For any intermediate cell state *i* and terminal state *j*, we can integrate over all possible paths (weighted by their probability) between *i* and *j*, thus computing the probability a cell starting at *i* will terminate at *j*. For each cell *i*, we can compute a vector of *branch probabilities* to reach each of the possible terminal states (Supp. Fig. 1f,g). We define its differentiation potential to be the entropy over the branch probabilities, providing a novel quantitative metric for its plasticity (Fig. 1d, Supp. Fig 1h).

Palantir assigns each cell both a pseudo-time (representing its relative distance from the start) and branch probabilities to *all* of the terminal states. Thus, unlike using diffusion components to order cells (where the order of cells changes across components), Palantir’s pseudo-time provides a single *unified* ordering of cells, across all lineages. This unified framework enables precise alignment, characterization and comparison of gene expression dynamics along different lineages, without having to sub-select cells of one or many lineages (Methods). From this pseudo-time ordering we compute gene expression trends using generalized additive models (GAMs), weighing each cell’s contribution based on branch probabilities (Fig 1e, Supp. Fig. 3, Methods). GAMs are particularly suitable for deriving a robust estimate of non-linear trends and estimating the standard error of prediction ^25^.

### Landscape of early human hematopoiesis

Hematopoiesis is a well-studied biological process with established markers to facilitate identification of lineages and terminal cell states ^17, 26^. Moreover, hematopoietic differentiation is asynchronous with the full spectrum of cell states represented in a single bone marrow sample. For these reasons, many pseudo-time algorithms have been developed using hematopoiesis as a model system ^2, 7, 9^. While scRNA-seq has been extensively used to study hematopoiesis in mouse ^6, 27^, we chose to investigate the earliest stages of human hematopoiesis, since single cell studies are particularly empowering in a system where perturbations are not possible.

Hematopoiesis has classically been characterized as a series of bifurcations leading to mature, terminal cell states ^17, 28, 29^. However, recent studies using scRNA-seq and scATAC-seq profiling of sorted populations suggest fate decisions in hematopoiesis is a continuous process ^4, 5^, raising a debate regarding the discrete versus continuous nature of differentiation in hematopoiesis. Despite the depth of our understanding of hematopoiesis, fundamental questions about how cell fate choice is determined at the earliest stages of human hematopoiesis, and the degree of plasticity in early progenitor cells, remain unanswered. To better investigate cell fate choices in early human hematopoiesis, we enriched for CD34+ cells using Fluorescent-Activated Cell Sorting (FACS) and generated approximately 25,000 single cell transcriptome profiles of CD34+ cells from three human bone marrow donors using 10X Chromium (Methods).

We first clustered the scRNA-seq profiles (Supp. Fig. 4a) using PhenoGraph ^20^. To associate each cluster with a particular cell type / lineage we used correlation between cluster medians and bulk sorted populations ^30, 31^ (Supp. Fig. 4b,c). We identified the full complement of hematopoietic cells, including hematopoietic stem and progenitors, as well as cells committed to lymphoid, erythroid, monocytic, classical and plasmacytoid dendritic cell (cDCs % pDCs respectively) lineages and megakaryocytes (Fig. 2a-b, Supp. Fig. 4). Stem and precursor cells comprised ~63% of the total sorted cells. Lineage committed cells were also detected because of imperfect sorting for CD34 (which was measured to be 90% pure) and temporal discordance between mRNA and protein; surface protein levels lag changes in mRNA levels.

**Fig 2:**
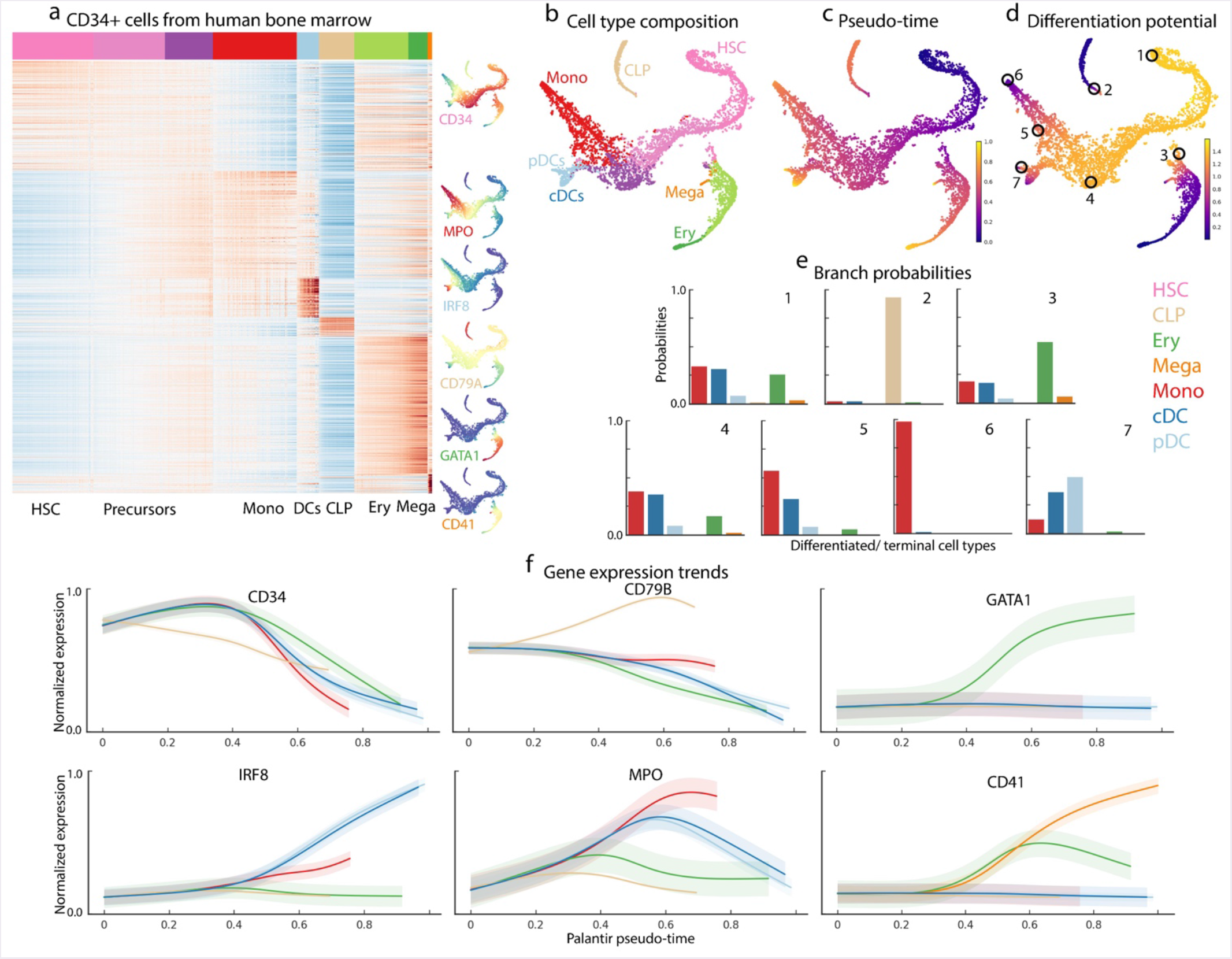
Differentiation landscape of early human hematopoiesis using CD34+ cells. Data shown for CD34+ cells from human bone marrow, replicate 1 (a) Heatmap showing the differentially expressed genes (based on MAST ^65^) between PhenoGraph ^20^ clusters. Each row represents a gene and each column represents a cell. Cells are ordered by cluster (listed at the bottom), top row represents cluster label, color coded in a consistent scheme throughout the rest of the figures and annotated on the bottom. tSNE maps show expression patterns for characteristic markers of different cell lineages. In each map, cells are colored based on expression of the labeled gene, blue for low expression and red for high expression. (b-d) tSNE maps of full scRNA-seq human hematopoiesis dataset, replicate 1. The results were generated using one of the HSCs as a start cell and the terminal cells were determined automatically. (b) Cells colored based on cluster label, color coded consistently with (a) and cell-type annotation is determined through correlation with bulk sorted populations. (c) Cells colored by Palantir pseudo-time and (d) Cells colored by Palantir differentiation potential (DP) (e) Histogram representation of branch probabilities of example cells highlighted in (d), each bar is color coded based on (b): (1) Early cells are capable of differentiating to all lineages, reflected as non-zero branch probabilities for all lineages. (2-3) lymphoid and erythroid lineages (2-3). monocyte and DC lineages (4-7). Note the gradual change in branch probabilities indicating continuity in cell fate. (f) Gene expression trends for characteristic genes for the different lineages, plotted as illustrated in Supp. Fig 3: CD34, a marker of stem and precursor cells is downregulated across all lineages; MPO, a myeloid marker shows an initial upregulation in all lineages and is downregulated in non-myeloid lineages; IRF8, CD79A and GATA1, lineage markers for DCs, CLPs and erythroid cells are upregulated in specifically in the corresponding lineages and CD41, a megakaryocyte marker shows an initial upregulated in the erythroid progenitors followed by a upregulation in the megakaryocytic lineage cells.

### Application of Palantir to hematopoiesis data recapitulates expected trends

We applied Palantir to the CD34+ hematopoiesis data, selecting one of the CD34 high cells as the start cell (Methods). We analyzed each of the three replicates separately to evaluate the robustness and reproducibility of the results. Based on the stationary distribution of the Markov chain, Palantir correctly identified all expected cell types, including: monocytes, erythroid cells, megakaryocytes, lymphoid progenitors and the two dendritic cell populations as terminal states. (Fig. 2b,c). The trajectory identified by Palantir begins at the HSCs and ends at the terminal differentiated cell types, following the expected progression (Fig. 2c). We observed a large degree of plasticity: cells at the beginning of the trajectory have potential to reach the full set of terminal states, with a gradual loss in this potential as they commit towards a particular lineage (Fig. 2d-e).

To evaluate the resulting trajectories, we computed gene expression trends of key markers that characterize commitment towards specific lineages (Fig. 2f). As expected, CD34 shows a decreasing trend in all lineages consistent with previous reports of CD34 downregulation as cells commit towards specific lineages ^17^. Consistent with well characterized trends, lineage specific factors such as CD79A, GATA1 and IRF8 are selectively upregulated in the lymphoid, erythroid and DC cell lineages respectively. In contrast to these genes, MPO shows an initial upward trend across all lineages, followed by a specific upregulation in the monocyte lineage (Fig. 2f). Finally, CD41 has been identified as a marker for early erythroid and megakaryocytic precursors with a continued upregulation in the megakaryocytic lineages ^32^.

We next evaluated Palantir’s robustness and reproducibility. Our experiments demonstrate that both pseudo-time and differentiation potential (DP) are robust to a wide range of input parameters including the number of neighbors for graph construction, number of diffusion components and different sampling of waypoints (Supp. Fig. 5, Methods). We also compared Palantir results between independent replicates (Methods) and observe that the pseudo-time and DP are highly correlated between independent runs of Palantir, each applied to datasets derived from different bone marrow donors. (Supp. Figs 6, 7, 8). Additionally, the gene expression trends are reproducible across the replicates (Supp. Fig. 7a). These findings collectively suggest that Palantir results are robust, reproducible, and correctly identify the gene expression dynamics in early hematopoiesis.

### Hematopoiesis data supports a hierarchical and continuous model of cell fate choice

Classically, hematopoiesis has been thought be a series of bifurcations through well-defined stages ^17, 28, 29^. However, these results were largely based on either a small number of markers or bulk measurements of sorted populations ^17, 31^. Recent studies point to a consensus supporting a more continuous model of hematopoiesis but disagree on whether differentiation proceeds in a hierarchical or non-hierarchical manner. On the one hand, a number of single-cell studies ^4, 6^ have hypothesized that hematopoietic decision making process is continuous but lacks hierarchy. These studies however were based on sorted populations and hence might have missed intermediate cell stages and more importantly did not retain the relative proportions of the different cell types. On the other hand, lineage-tracing studies of murine hematopoiesis ^33^ support a hierarchical model of development with a more stepwise loss in potential as stem cells differentiate into specific cell types.

Our Palantir framework and the CD34+ data allow us to query early differentiation in *human* hematopoiesis where genetic perturbation studies are impossible. Key to our approach is comparing the change in differentiation potential (DP), across different lineages. DP shows a decreasing trend along any given lineage, as cells specialize and lose their ability to commit to other lineages (Supp. Fig. 9a-d). Tracking branch probabilities (BPs) and DP along pseudo-time enables us to determine when and in what manner these probabilities change for each of the terminal fates. Our results suggest continuity in the early hematopoietic lineage commitment process: DP remains consistently high throughout early hematopoiesis, with gradual losses in potential as cells differentiate towards specific lineages (Fig. 3a, Supp. Fig. 9e).

**Fig 3:**
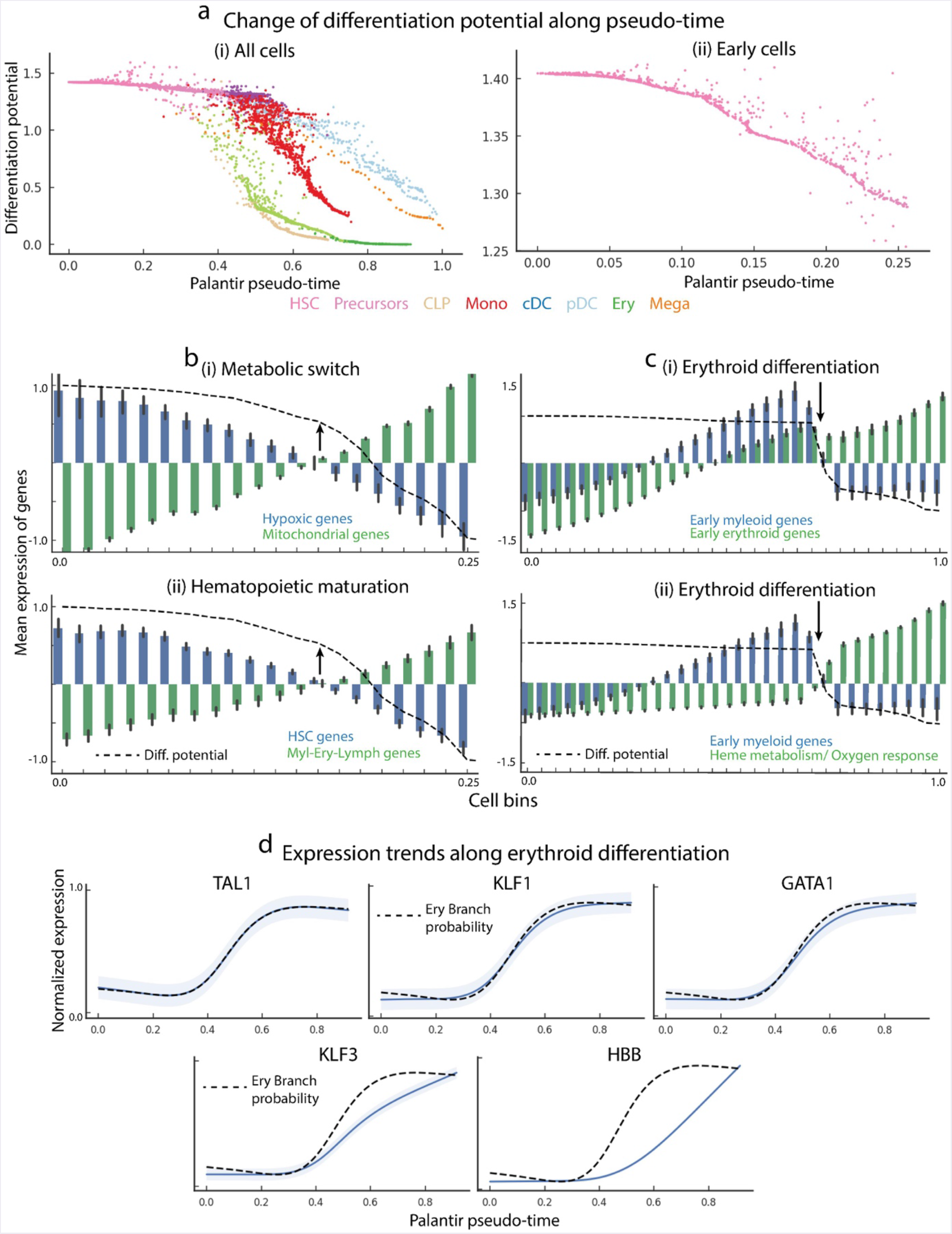
Palantir differentiation potential identifies landmarks of hematopoietic differentiation. Data represents results for CD34+ cells from human bone marrow, replicate 1, as in Figure 2. (a) Plots showing differentiation potential along pseudo-time. (i) Each dot represents a cell, color coded based on clusters in Fig 2b. X-axis is pseudo-time and Y-axis is DP. DP shows a decreasing trend as cells commit to different lineages. (ii) Similar plots, zooming into the early hematopoietic cells. (b) (i) Plot showing the average expression (y-axis, left) for early hematopoietic cells of hypoxic genes (blue) and mitochondrial genes (green) in equal sized bins along Palantir pseudo-time (x-axis). Dotted black line represents DP (y-axis, right), plotted along the corresponding pseudotime. Arrow indicates the point of maximal DP change. DP drop corresponds to the region where expression of mitochondrial genes exceeds the expression of hypoxic genes. (ii) Similar to top panel, comparing the expression of stem cell genes (blue) and mature cell lineage specifying genes (green), with DP shown in black. (c) Similar to (b), plots showing the expression of different sets of genes, binning the cells along erythroid lineage. (i) Plots showing average expression of early myeloid genes (blue) and early erythroid genes (green). DP in each bin is shown in black. The point of maximal DP change correlates with higher expression of erythroid genes. (ii) Similar to top panel, with the genes involved in functional specification of erythroid function shown in in green. (d) Top panel: Gene expression trends (see methods for computation) of key erythroid TFs: TAL1, KLF1 and GATA1 which are the most correlated with erythroid branch probability (shown in black). Trends are plotted in blue with shaded region representing 1 standard deviation. Bottom panel: Gene expression trends of KLF3 and HBB.

Importantly, we note that the *rate of change* in DP varies greatly along pseudo-time and across the different lineages (Fig 3a, Supp. Fig. 9e, Methods). If lineage commitment was non-hierarchical, we would expect DP along different lineages to simultaneously drop downward at a particular point along pseudo-time. Instead, we observe first commitment to the lymphoid lineage, followed by commitment to erythroid/megakaryocytic lineages and finally we observe cells differentiating towards the myeloid lineages (Fig 3a, Supp. Fig. 9e), supporting a hierarchical nature of human hematopoietic lineage commitment. Our results also suggest that lineage choice between either the monocytic or one of the two DC lineages precedes fate choice between the two DC lineages (Supp. Fig. 9e). Together, these results suggest that differentiation in early human hematopoiesis exhibits hierarchy.

### Differentiation potential identifies landmarks of early hematopoietic differentiation

During differentiation, cells lose their fullest multipotent potential as they commit towards particular lineages. Differentiation potential (DP) represents a quantitative measure of a cell’s plasticity or potential to differentiate into different lineages. DP can thus detect when cell fate specification changes. We observe points along the differentiation axis (pseudo-time) where substantial changes in DP occur and posit that these changes in DP reflect key molecular and cellular events driving differentiation. Indeed, most of these changes coincide with commitment to different lineages (Fig. 3a left panel, Supp. Fig. 9), Curiously, we observe a substantial decrease in DP in early hematopoietic differentiation (Fig 3a top right panel) not associated with commitment towards any specific lineage.

The DP metric reflects the graph structure, which in turn is constructed based on genome-wide similarities between cells. Hence to gain insight into this drop in DP, we sought to characterize gene expression trends in proximity to this event. We clustered genes based upon their temporal trends along the pseudo-time, assuming genes involved in coherent biological processes likely share similar expression dynamics and subsequently used gene ontology enrichment to annotate the resulting clusters (Methods).

The strongest changes we observed involved: (1) aerobic and mitochondrial respiration related genes (clusters with increasing trends) and conversely (2) hypoxic genes (clusters with decreasing trends) (Supp. Fig. 10a). To better visualize and quantify the correspondence between these gene trends and DP, we compared the mean expression of hypoxic genes with genes involved in mitochondrial respiration along equal sized bins (Fig. 3b, top panel). As shown in Fig. 3b, decrease in DP strongly correlates with both an increased expression of mitochondrial genes and a decreased expression of hypoxic genes. These data suggest a decrease in DP at the earliest stages of hematopoiesis corresponds with a change in metabolic state of the cells, occurring before they begin to commit towards lineages (Fig. 3b, top panel).

Previous studies have shown that hematopoietic stem cells (HSCs) are inherently quiescent, slow cycling in nature, and reside in specialized hypoxic niches in the bone marrow ^34, 35^. Hematopoietic differentiation requires an exit from quiescent long-term HSCs (LT-HSCs) to a metabolically active short-term HSCs (ST-HSCs), a process termed the metabolic switch ^34, 35^. The complement of cell types into which a cell can differentiate is thought to remain unaltered during the transition of LT-HSCs to ST-HSCs. Consistent with these studies, we show that the change in DP correlates with the metabolic switch reproducibly and independently in each of the three replicate samples (Fig. 3b, Supp. Fig. 11). DP change is also correlated with expression dynamics of THY1(CD90), a well characterized marker of transition between LT-HSCs to ST-HSCs (Supp. Fig. 10b) ^36^. Moreover, change in DP is also accompanied by increased expression of early myeloid-erythroid-lymphoid genes in comparison to genes that are characteristic of hematopoietic stem cells (Fig. 3b, Supp. Fig. 11). These results demonstrate that DP, as computed by Palantir directly from the data with no use of prior knowledge, can identify key differentiation events, such as metabolic switch, even when these are unrelated to specific branching in cell fate.

### Differentiation potential along erythroid commitment

We next characterized DP changes along lineage commitment using erythropoiesis, a process that produces red blood cells as a case study. Erythrocytes are derived from megakaryocyte-erythroid precursors (MEPs), lineage precursors of both megakaryocytes and erythrocytes ^37^. We observe a sharp decrease in DP along erythroid commitment after the initial, metabolically-related entropy change during early hematopoiesis (Fig 3a). To identify gene sets concordant with this decrease in DP along erythroid differentiation, we repeated the temporal trend-based gene set analysis as before (Supp. Fig. 10c).

During early hematopoiesis, we observe an increased expression of gene markers across all differentiated cell types (Fig 3b). Gene expression trends in cells demonstrating commitment towards erythroid lineage (increasing BP toward erythroid cell fate) are associated with continued upregulation of early erythroid genes, accompanied by a downregulation of the early myeloid genes (Fig. 3c), demonstrating a reconfiguration of the expression program toward erythroid lineage. As expected for maturing red blood cells, this decrease in DP also coincides with upregulation of gene sets involved in heme metabolism and oxygen response (Fig. 3c).

We posited that the transcription factors (TFs) most closely correlated with the BP towards the erythroid fate are likely key regulators of this process. Hence, we systematically correlated all TFs with erythroid BP and found the most correlated TFs to be TAL1, KLF1 and GATA1 (Pearson correlation > 0.99) (Fig 3d, Supp. Fig. 10d (Cluster 0)). We note that each of these factors have been shown to play a central role in erythropoiesis. TAL1 is known to enhance erythroid potential ^38^. KLF1 is a known regulator of early erythroid precursor genes, as well as a suppressor of the megakaryocyte lineage ^39^. Finally, loss of GATA1 leads to complete loss of erythropoiesis ^40^. Thus, we find remarkable correspondence between erythroid BP, computed based on all genes with no prior knowledge, and expression trends of known key regulators of erythropoiesis.

Palantir provides a high resolution cell ordering along differentiation, allowing us to characterize the order and timing of events along erythropoiesis. We find that upregulation of KLF1 is followed by upregulation of KLF3, a known target of KLF1 and a stabilizer of the erythroid gene expression program (Fig. 3d, middle panel, Supp. Fig. 10d (Cluster 6)) ^41^. Finally, globin genes such as HBB are upregulated in the final wave conferring functional identity to red blood cells (Fig. 3d, right pane, Supp. Fig. 10d (Cluster 8)). Taken together, the timing of gene upregulation along the erythropoietic trajectory strongly suggests that specification and commitment to erythroid lineage occurs not equipotentially, but in stages of coordinated upregulation of genes.

### Transcriptional regulation of erythroid commitment

Given the strong correspondence between key erythroid TF expression and erythroid BP, we next sought to use Palantir to identify factors that influence lineage fate choices. We reasoned that a TF that influences lineage decisions should satisfy the following criteria: (a) The factor should be expressed prior to phenotypic specification i.e., before the lineage decision is made, (b) The factor should correlate with lineage specification i.e., upregulation of the factor during the early phase of specification should be correlated with increasing lineage probability and (c) The factor should be downregulated in alternate lineages.

Considering the specific case of myeloid-versus-erythroid lineage decision, mutual antagonism of GATA1 and PU.1 has been proposed as the primary mechanism. Specifically, PU.1 and GATA1 drive myeloid and erythroid specification respectively ^42, 43^. Our data supports PU.1 as a myeloid specifier: (a) PU.1 is expressed in early hematopoietic stem cells, (b) PU.1 shows an increasing trend along myeloid lineages and (c) PU.1 is downregulated in the erythroid lineage (Fig. 4a, Supp. Fig. 12a). On the other hand, GATA1 expression is not detected in early cells and is only expressed in the erythroid lineage after the cell fate decision has been made (Fig. 4a). Therefore, it is unlikely that GATA1 plays a role in the erythroid fate decision, a finding consistent with recent studies based on time-lapse microscopy ^44^. Hence, we sought to discover alternative factors that could antagonize PU.1 and drive cells towards the erythroid lineage.

**Fig 4:**
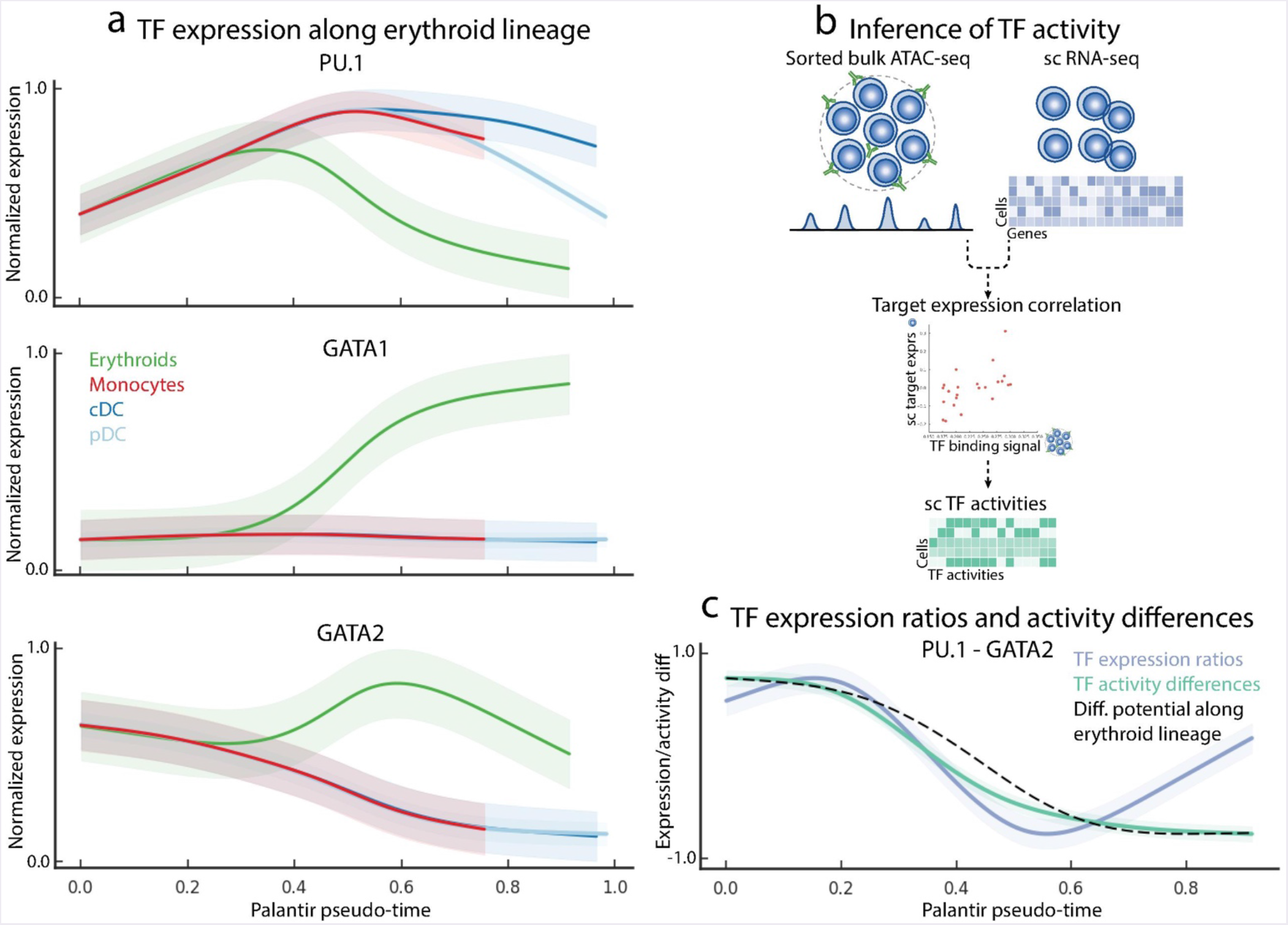
Transcriptional regulation of erythroid differentiation. Data represents Palantir results for CD34+ cells from human bone marrow, replicate 1. (a) Gene expression trends for PU.1, GATA1 and GATA2 in the myeloid and erythroid lineages. The trends are color coded based on Fig. 2b. (b) Schematic representation of single cell TF activity inference using scRNA-seq data and ATAC-seq data from bulk sorted populations. ATAC-seq data is used to identify cell-type specific TF targets and TF activity in each cell is inferred by measuring the correlation between predicted TF sequence affinity of the targets with their expression. (c) Trend plots showing the dynamics of PU.1 - GATA2 expression ratio (blue), PU.1 - GATA2 TF activity difference (green) along the erythroid lineage. DP change along the same lineage is shown in black. Change in PU.1 - GATA2 expression ratios and TF activity differences strongly correlate with DP change along erythroid lineage.

To do so, we undertook a systematic evaluation of all the TFs expressed in the erythroid lineage that satisfy the criteria articulated above. For each TF, we computed the correlation of its expression trend with the increase in erythroid BP during lineage specification and also considered its expression in the early hematopoietic cells (Methods). We found that GATA2, LYL1 and MXD4 all exhibit high correlation with erythroid commitment and are expressed at high levels in the precursor cells (Supp. Fig. 12b). GATA2 has the highest correlation and is the highest expressed amongst these factors. Moreover, GATA2 belongs to the same TF family as GATA1 and two factors share the same binding motif. We note that GATA1 and PU.1 are thought to drive respective lineage commitment by an antagonistic mechanism of binding to each other’s promoters ^42, 43^ and hence GATA2 could function as a PU.1 agonist using the same mechanism. In fact, switching of GATA factors is a well characterized mechanism during erythroid differentiation and our data again supports this: GATA2 is upregulated earlier than GATA1 and then decreases along with GATA1 upregulation (Fig. 4a). Thus, our high resolution temporal model suggests that GATA2 drives erythroid lineage specification by antagonizing PU.1, with GATA1 taking over from GATA2 during lineage commitment. We therefore chose GATA2 as a candidate PU.1 agonist and erythroid lineage specifier.

We leveraged changes in the DP to investigate the timing of the decrease in cell plasticity and its interplay with the changes in TF expression driving lineage specification. Previous studies have shown that gene expression *ratios* between competing TF pairs, rather than TF expression alone, can be critical determinants of lineage specification ^45^. TF ratios can specify different lineages by either directly repressing the factor in the pair or by activating a repressor of the other factor ^42, 43, 46^. While average GATA2 levels remain relatively constant during early hematopoiesis (Supp. Fig. 12c), the multi-dimensional nature of our single-cell data enables us to examine how ratios between TF pairs might change during the course of differentiation. Indeed, we observe that the decrease in ratio of PU.1/GATA2 expression precedes the drop in DP (Supp. Fig. 12d). This suggests that as the balance of expression between PU.1 and GATA2 tilts towards GATA2 dominance, regulatory mechanisms initiate gene expression programs that confer erythroid fate and indeed the expression ratio of PU.1/GATA2 is correlated with DP change along the erythroid lineage (Fig. 4c, blue line).

To gain a better understanding of the downstream impacts of the PU.1/GATA2 ratio (and DP change), we set out to characterize the behavior of PU.1 and GATA2 *target genes* along the erythroid lineage. Concordant behavior of multiple target genes not only mitigates individual gene measurement noise in scRNA-seq, but also provides a functional readout of the TF activity. We note that the targets of a TF largely depend on the tissue or cell type. Therefore, we leveraged previously published bulk ATAC-seq data ^12^ from sorted Erythroid cells for GATA2 targets and GMP cells for PU.1 targets to determine TF activities at single cell level (Fig. 4b, Methods). The expression of TF targets has been shown to be strongly correlated with sequence affinity of the TF to its targets ^11^. Therefore, we identified TF binding sites and targets from ATAC-seq data ^47^ and computed the correlation between predicted sequence affinity and target expression to infer TF activities for each factor and cell (Supp. Fig. 12e, Methods). In line with the expression ratios, we observe that the change in PU.1 and GATA activity difference precedes the change in DP (Supp. Fig. 12d). We note that the activity difference is also strongly correlated with the decrease in DP along the erythroid lineage (Fig. 4c, green line). Our findings are independently reproducible across the three replicates (Supp. Fig. 12f-g). Together, these results provide further evidence that GATA2, rather than GATA1, functions as a mutual agonist of PU.1 to achieve erythroid specification.

### Application of Palantir to mouse hematopoiesis and colon differentiation

Palantir is ideally suited for our CD34+ human hematopoiesis dataset, which was heavily enriched for early multipotent precursors, providing sufficient early cells for fine resolution mapping of lineage fate decisions. We wanted to test Palantir in a more challenging setting, where there is a paucity of early cells and potential bias induced by cell sorting. We therefore selected a mouse hematopoiesis dataset that profiled Lin^-^c-Kit^+^Sca-1^+^ cells using MARS-seq2 ^6^. This study specifically sorted cells for different precursor populations (monocytes, neutrophils, basophils, DCs, erythrocytes and megakaryocytes), but excluded the most multipotent stem cells, thus creating a challenge to correctly resolve branching probabilities (note paucity of early cells in Fig. 5a). In addition to functioning as a test case, this dataset also allows a qualitative comparison between human and mouse hematopoiesis.

**Fig 5:**
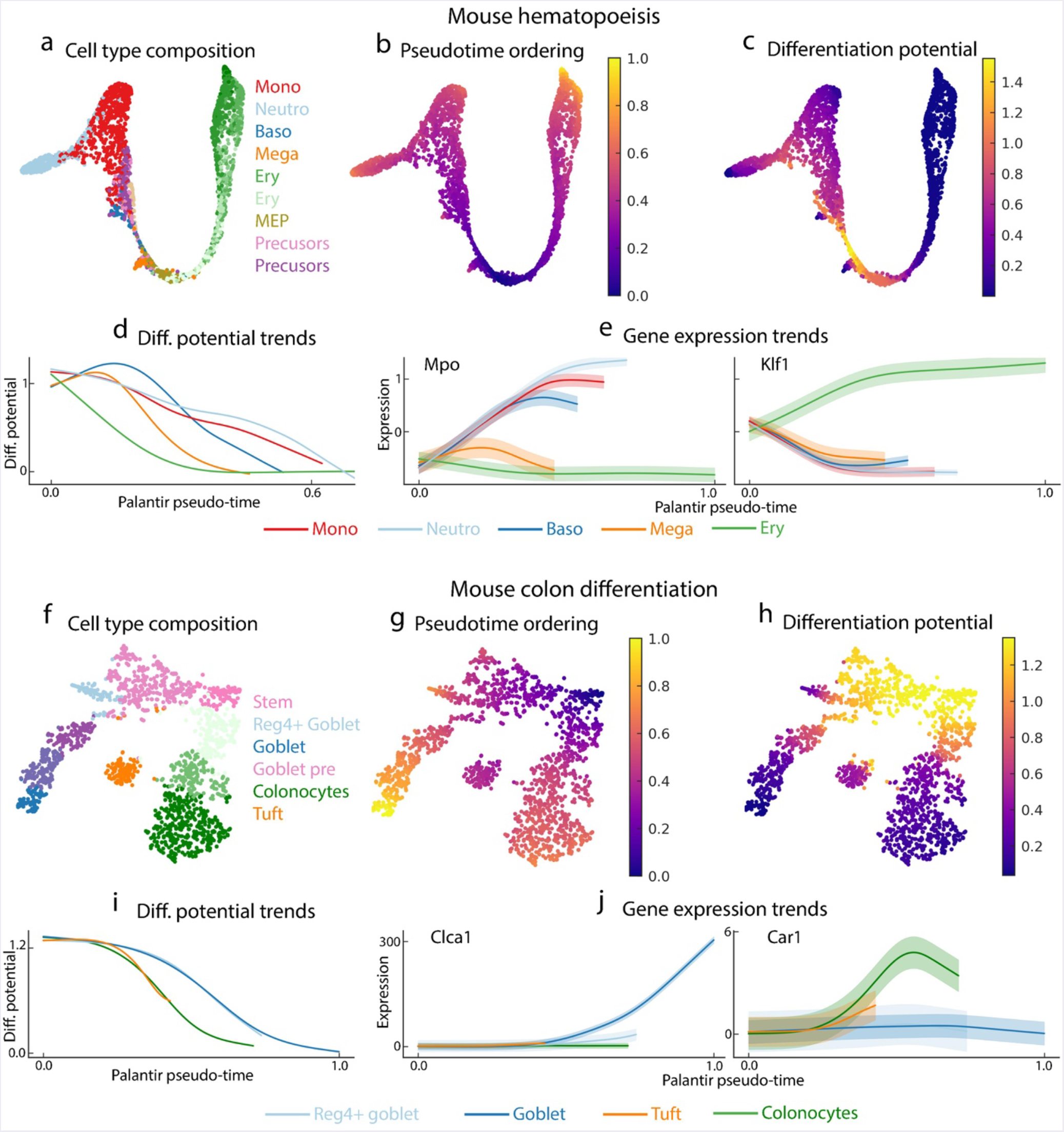
Palantir generalizes to mouse hematopoiesis and colon differentiation datasets. (a) tSNE map of mouse hematopoiesis data generated by scRNA-seq of sorted precursor populations ^6^. Cell are color coded based on clusters generated by ^6^. Notice the paucity of well-defined stem cell population. (b-c) Cells colored by Palantir pseudo-time and differentiation potential generated using an early cell as the start. (d) Plot showing the DP trends along pseudo-time for the different lineages. Trends are color coded based on clusters in Fig 5a. The changes highlight the hierarchical nature of murine hematopoiesis with commitment towards the erythroid lineage (green) followed by commitment towards the different myeloid lineages. (e) Gene expression trends of Mpo and Klf1, myeloid and erythroid factors respectively, recapitulate their expected behavior and are consistent with their dynamics in human hematopoiesis. (f) tSNE map of scRNA-seq dataset of epithelial enriched cells from the mouse colon ^48^. Cells are color coded based on Phenograph clusters. (g-h) Cells colored Palantir pseudo-time and DP generated using one of the Lgr5+ stem cells as the start cell. Results were generated by manually setting the tuft cells as one of the terminal states. (i) Plot showing the DP trends for all the lineages. Trends are color coded based on clusters in Fig. 5f. The DP trends recapitulate the known hierarchy of lineage specification with specification of colonocytes (green) following by differentiation towards the two goblet cell populations (blue). (j) Gene expression trends of Clca1 and Car1 across different lineages.

Provided only with few multipotent precursor cells, Palantir was still able to correctly identify the different terminal states and estimate pseudo-time and DP characterizing mouse hematopoiesis (Fig. 5b-c, Supp. Fig. 13a, Methods). We observe that the peak DP is not at the beginning (Fig. 5d) and attribute this to the paucity of multipotent cell populations - which affects the accuracy and resolution of the multipotent cells at the beginning. Despite these limitations, we observe a clear hierarchical structure in lineage specification (Fig. 5d), consistent with recent lineage tracing experiments ^33^. The hierarchical structure is similar to human hematopoiesis, with commitment to erythroid lineage followed by specification of the different myeloid lineages (Fig. 5d). As further support of Palantir’s model, the expression trends of a key erythroid and myeloid genes, Mpo and Klf1 are consistent with their role in respective lineages (Fig. 5e) ^39^ and follow similar patterns to their behavior in human hematopoiesis (Fig. 2f, 3d), demonstrating the robustness of Palantir to altered composition of differentiating cell types.

To test the generalization of Palantir to non-hematopoietic datasets, we applied Palantir to scRNA-seq dataset of mouse colon differentiation generated using the InDrop platform ^48^. Lgr5+ stem cells were shown to differentiate to colonocytes, tuft cells, goblet cells and Reg4+ goblet cells (Fig. 5f). Palantir automatically identified the two goblet populations and colonocytes as terminal states but failed to identify Tuft cells as a terminal state. Tuft cells were not identified as a separate terminal state since this population of cells is not completely mature and closer to Lgr5+ cells compared to the other terminal cell types. (Fig. 5f-g). By manually setting Tuft cells as one of the terminal states, Palantir correctly identified the pseudo-time ordering, hierarchical relationships and order of lineage commitment in mouse colon differentiation: lineage specification of colonocytes is followed by lineage specification towards the goblet cell populations (Fig. 5g-i, Supp. Fig 13b) ^49^. Additionally, gene expression trends recapitulate expected behaviors along different lineages: Clca1 is specifically upregulated in the goblet cell lineage; Car1 shows an initial upregulation in the colonocytes followed by a marginal downregulation, accurately capturing the behavior of these genes along the respective lineages; Muc1 shows stronger upregulation in Reg4+ goblet cells and Lgr5, the stem cell marker shows a downward trend across all lineages (Fig. 5j, Supp. Fig. 13c) ^48^. To evaluate Palantir’s robustness to missing terminal states, we also ran Palantir without manual intervention, i.e. when Tuft cells were not used as one of the terminal states. The BP changes and expression trends along lineages other than Tuft were not significantly altered (Correlation: 0.98; Supp. Fig. 13d), demonstrating that Palantir is robust to missing populations and mislabeled cells. These results demonstrate that continuities in cell fate choices are widespread across various biological systems and Palantir is uniquely positioned to model these continuities and thus enables characterization of lineage decisions across diverse datasets, including correctly recapitulating the dynamics of key TFs that regulate these processes.

### Comparison of Palantir to other pseudotime algorithms

Palantir is designed to investigate cell plasticity and fate decisions, based upon a continuous, probabilistic model for a cell’s potential to reach different cell fates. While significant advances have been made for resolving pseudo-time ordering of cells, state of the art pseudo-time algorithms continue to model differentiation as a series of discrete, deterministic bifurcations, predominantly approximated by clustering the data ^7, 8^. To benchmark its performance and outputs, we compared Palantir to leading and widely used pseudo-time algorithms such as Slingshot^8^, Partition Based Graph Abstraction ^7^, Monocle^9^, and diffusion maps ^3^. We used our human hematopoiesis data to compare the different algorithms and evaluate their ability to characterize a complex differentiating system.

Diffusion maps are widely applied for pseudo-time ordering of cells by projecting cells along *individual* components to determine pseudo-time for a lineage. Gene expression trends are then estimated by using a sliding window approach along these projections ^23, 24^. There are three key limitations to this approach: First, projection of cells onto a *single* diffusion component does not always generate an accurate ordering of cells along a lineage. Second, diffusion maps generate a projection of *all* cells along each component and therefore segmentation of the data (e.g. based on clustering) is necessary to determine gene expression dynamics. Finally, projection of cells along different components does not allow for a *direct comparison* of dynamics between two different lineages.

In particular, our data demonstrate that only the monocytic and lymphoid lineages can be unambiguously explained by a single diffusion component; all other lineages require multiple components to accurately determine pseudo-time (Supp. Fig. 2b). Bearing these limits in mind, we used projection of lymphoid lineage cells to characterize the effect of using individual diffusion components for determining pseudo-time and gene expression trends (Supp. Fig. 14a(i)). Projections along this component amplify the already-significant density differences in the data (Supp. Fig. 14a(i)). Sliding window approaches are particularly sensitive to density differences and as a result, the expression trend estimates are not reliable. In this particular case, loss of resolution in the sliding window approach prevents an accurate characterization of key TFs such as PU.1 (Supp. Fig. 14a(ii)), which has been shown to play a key role in lymphoid specification ^50^.

We next compared our results to Monocle2, a widely used trajectory detection algorithm ^9^, which uses reverse graph embedding to simultaneously learn the principle curve explaining the manifold of the data and projections of cells onto the curves. Monocle2 applied to human hematopoiesis using default parameters identified six distinct states in the data, but we could not attribute specific cell types to any of these states based on expression of marker genes (Supp. Fig. 14b(i-ii)). Moreover, key canonical markers for progenitors (CD34), myeloid (MPO, IRF8) and B-cell lineages (CD79B) are spread across all projections with no trend or coherence. Thus, Monocole2 failed to correctly compute pseudo-time, identify terminal fates and generate expression trends on this data. We note that in the original publication, Monocle2 was demonstrated on a small dataset, with well distinguished, sorted populations, rather than a complex differentiating system.

We next applied Partition Based Graph Abstraction (PAGA) ^7^ to the human hematopoiesis data. PAGA aims to reconcile pseudo-time by approximate graph abstraction and is particularly adept at characterizing large datasets with different complexities ^7^. PAGA partially succeeds in recovering the different hematopoietic lineages and their relationships (Supp. Fig. 14c). However, PAGA embeds the megakaryocyte lineage cells into the erythroid cell lineage and is unable to distinguish between the two DC lineages (Supp. Fig. 14c(ii)). The abstracted graph constructed by PAGA represents the lineage decision process, thus the strong interconnectivity among clusters representing the intermediate states provides further evidence for lack of well-defined bifurcations in human hematopoiesis (Supp. Fig. 14c(ii)). PAGA uses a sliding window to infer gene expression trends and requires a manual specification of clusters that contribute to a particular lineage. As demonstrated above, the sliding window approaches are sensitive to density differences in the data and do not generate sufficiently reliable estimates to characterize key events along lineage commitment (Supp. Fig. 14c(iii)).

Next, we applied Slingshot ^8^, an algorithm that clusters the data and constructs a minimum spanning tree (MST) through the clusters to identify the different lineages and pseudo-time. Slingshot does not make explicit recommendations for dimensionality reduction and clustering algorithms, both required as input. Therefore, to maximize similarity to Palantir, we applied Slingshot to the hematopoiesis data using diffusion maps ^21^ and Phenograph ^20^ clusters as input and recovered four lineages from the data (monocyte, lymphoid, erythroid and DC) with (Supp. Fig. 14d(i)). Similar to PAGA, Slingshot fails to distinguish between the two DC clusters (since they are clustered together) and embeds megakaryocyte population to be a stage along erythroid lineage, even though these are clustered separately (Supp. Fig. 14d(i), Supp. Fig 4a). Slingshot relies centrally on clustering and consequently cells committing towards Myeloid lineage are included as part of the lymphoid lineage (Supp. Fig. 14d(i) - Lineage 2). This results in an unexpected downward trend in CD79B along this lineage (Supp. Fig. 14d(ii)). Since the gene expression trends for the two DC lineages are identical, we cannot distinguish between expression dynamics of key DC TFs such as CEBPG (Supp. Fig. 14d(ii)).

A recently published approach, population balance analysis (PBA) ^14, 15^ presents a framework to characterize differentiation using spectral graph theory to solve a system of differential equations representing the dynamics of maturation along a lineage. In practice, this translates to using Markov chains for characterizing differentiation, providing further support for this approach to model differentiation. We note that PBA requires extensive use of prior knowledge to infer the proliferation and loss rates for each state (cell) in the system, which form the fundamental basis for the Markov chain construction. In the particular case of mouse hematopoiesis, the rates for the different lineages were estimated separately using data from multiple fate mapping studies ^14^. In addition, PBA requires explicit specification of the terminal states in the system *a priori*. We could not apply PBA to human hematopoiesis owing to paucity of such fate mapping studies in human. In contrast, Palantir requires only specification of an early cell and can automatically construct the Markov chain and determine the set of terminal states based on single cell RNA-seq measurements alone. This unbiased approach to characterizing differentiating systems is a key strength of Palantir’s utility and applicability to model tissue systems without established lineages.

We find that Palantir substantially outperforms the other algorithms in identifying the terminal states and recapitulating gene expression trends along differentiation. Importantly, none of the algorithms discussed above explicitly model and quantify the plasticity and branch probabilities resulting from continuities in cell fate choices. Taken together, only Palantir could accurately associate expression changes in key transcription factors with changes in commitment to the lineages these regulate.

## Discussion

scRNA-seq datasets have provided strong support for a continuous model of differentiation and emerging evidence suggests cell fate choice is likewise continuous in nature. We have developed Palantir, the first algorithm to model these continuities by determining, for each cell state, branch probabilities for each lineage, and quantifying the differential potential along the differentiation landscape. Palantir is robust to different parameters, reproducible across replicates, and generalizes to diverse datasets. We applied Palantir to study human hematopoiesis, based on human bone-marrow data, enriched for stem and precursor populations. Palantir’s high resolution mapping of cells along distinct differentiation trajectories allowed us to characterize both the order and timing of key regulatory factors driving lineage choices. Our findings clarified cell fate choice in human hematopoiesis is a hierarchical process.

Key to the high resolution of Palantir pseudo-time is the use of multiple diffusion components and neighbor graphs to measure distances between cells in this embedded space (Supp. Fig. 15a-c). This high-resolution ordering enables Markov chain construction, which is central to both identifying terminal states and modeling continuities in lineage choices. Palantir’s unified framework of modeling continuities in cell state and fate choices allows us to identify key landmarks along hematopoiesis such as the metabolic switch in stem cells and gain insights into lineage decisions by comparing gene expression dynamics across lineages. Palantir outperforms other pseudo-time algorithms, which largely treat lineage choices as discrete bifurcations, in recovering gene expression trends and lineage relationships that are more consistent with known human hematopoiesis biology. The unbiased sorting of stem and precursor cells from bone marrow was critically important to characterize lineage choices in early human hematopoiesis at high resolution. However, Palantir can robustly recover expression trends in datasets where the precursors populations are not enriched.

While we have demonstrated the capabilities of Palantir in the well characterized hematopoiesis system, with the launch of the Human Cell Atlas Project ^51^, we anticipate that Palantir will be a valuable discovery tool for many less characterized systems. A key requisite for the success of Palantir is the presence of the full dynamic range of differentiating cells, made possible by the asynchronous nature of differentiation in certain tissues such as bone marrow, colon and olfactory epithelium ^8, 27, 48^. We note that this feature is not present in embryogenesis which is typically studied based on time course experiments ^24, 52, 53^. Characterization of the order and timing of events in such systems will require an explicit modeling of connectivity between the time points and correction of confounding factors introduced by measuring batches independently ^54^.

The most important assumption made by pseudo-time algorithms, including Palantir, is that differentiation is unidirectional from immature precursor cells to functionally mature cells. While this is a reasonable assumption in healthy differentiation, it has been demonstrated to be violated in systems such as tissue regeneration ^55^ and in cancer ^56^. However, if cells dedifferentiate or trans-differentiate to a state which is transcriptionally identical to an earlier state, scRNA-seq data alone is insufficient to characterize the differentiation paths. Recent advances in in-vivo lineage tracing technologies are starting to provide ground truth for lineage relationships ^57, 58^. However, these require genetically modified model systems and hence are unsuitable to study cancer progression, metastasis and healthy development in human tissues. As an alternative, one could use mutations, rapidly occurring in most cancers, to gain directionality and lineage information in the human system. Moreover, recent studies ^59^ have demonstrated that somatic mutations occur at a rate that allows for lineage tracing in healthy human tissues based on these mutations. The ability to simultaneously profile the transcriptome and DNA ^60^ has great potential to elucidate disease initiation and progression by extending Palantir to incorporate lineage information to model cell fate decisions.

### scRNA-seq of CD34+ human bone marrow cells

Cryopreserved bone marrow stem/progenitor CD34+ cells from healthy donors were purchased from AllCells, LLC. (Cat. No. ABM022F) and stored in vapor phase nitrogen until use. Typical for scRNA-seq, a vial was removed from the storage and immediately thawed at 37°C in a water bath for 2-3 minutes. Next, vial content (1ml) was transferred to a 50ml conical tube. In order to prevent osmotic lysis and ensure gradual loss of cryoprotectant, 1ml of warm media (IMDM with 10% FBS supplement) was added dropwise, while gently shaking the tube. Then the cell suspension was serially diluted 5 times with 1:1 volume additions of complete growth media with 2 minutes wait between additions. Final ~32ml volume of cell suspension was pelleted at 300rcf for 5 minutes. After removing supernatant, cells were washed twice in ice cold 1X PBS with 0.04% (wt/vol) BSA supplement to remove traces of media. Cell concentration and viability was determined with Countess II automatic cell counter employing trypan blue staining method.

Single cell RNA sequencing was performed with 10X genomics system using Chromium Single Cell 3` Library and Gel Bead Kit V2 (Cat. No. 120234). Briefly, 8700 cells (viability 90-97%) were loaded per reaction, targeting recovery of 5000 cells with 3.9% multiplet rate. After reverse transcription reaction emulsions were broken, barcoded cDNA was purified with DynaBeads, followed by 12 cycles of PCR amplification. The resulting amplified cDNA was sufficient to construct NGS libraries, which were sequenced on Illumina HiSeq 2500 system (HiSeq SBS V4 chemistry kit).

### Single cell RNA-seq data processing

#### Data preprocessing

Data derived from each replicate was processed independently. Single cell RNA-seq data was preprocessed using the SEQC pipeline ^30^ using hg38 human genome and the default SEQC parameters for 10X to obtain the molecule count matrix. The SEQC pipeline aligns the reads to the genome; corrects barcode and UMI errors; resolves multi-mapping reads and generates a molecule count matrix ^30^. SEQC also performs a number of filtering steps: (a) Identification of true cells from cumulative distribution of molecule counts per barcode, (b) removal of apoptotic cells identified at cells with >20% of molecules derived from the mitochondria and (c) removal of low complexity cells identified as cells where the detected molecules are aligned to a small subset of genes ^30^. In addition, cells with less than 1000 molecules detected were filtered out. Finally, genes that were detected in at least 10 cells were retained for downstream analysis.

The filtered count matrix was normalized by dividing the counts of each cell by the total molecule counts detected in that particular cell. The normalized matrix was multiplied by the median of total molecules across cells to avoid numerical issues ^61^. Normalized data was log transformed with a pseudocount of 0.1.

#### Cell cycle correction

Expression of cell cycle genes can confound the ordering of cells in a differentiation trajectory. and hence we applied f-scLVM ^62, 63^ to factor out the cell-cycle effect across all cells. Normalized and log transformed data was used as input to f-scLVM correction with default parameters. The following gene ontology annotations were used to annotate the cell cycle effect: GO:0000279 M phase, GO:0006260 DNA replication, GO:0007059 chromosome segregation, GO:0000087 M phase of mitotic cell cycle, GO:0048285 organelle fission.

Following cell cycle correction, PCA was performed keeping the top 300 components and diffusion maps were computed using the PCs as input ^21^. See section “Adaptive anisotropic kernel” under the Palantir algorithm description for details on constructing the diffusion maps.

#### Annotation of cell types and filtering of mature populations

Gene expression profiles from sorted bulk hematopoietic populations were used to annotate the cell types ^30, 31^. Cell cycle corrected data was clustered with Phenograph ^20^ using default parameters and the top 300 principle components as inputs. Cluster centroids were determined for each cluster and the expression of each gene was standardized. Bulk expression data was downloaded from the Dmap portal (http://portals.broadinstitute.org/dmap/home) and expression of each cell type was standardized. For each cluster, average correlation across bulk replicates was computed for each cell type and the cell type with the highest correlation was used to annotate the cluster (Supp. Fig. 4c). Note, the inferred cell types are used only for interpretation and not used by Palantir.

To limit the data to cell types undergoing differentiation in the bone marrow, clusters that were annotated as T-cells and mature granulocytes were filtered out. T cells were filtered out, since these migrate from the periphery and do not differentiate in the bone marrow. Mature granulocytes were filtered out since no coherent precursor population was identified in the data.

#### tSNE visualization

tSNE maps ^64^ were generated using diffusion components scaled by the Eigenvalues as inputs rather than principal components of the data and perplexity set to 150. The scaling of Eigenvectors ensures less sensitivity to outliers in the data and is performed as follow:

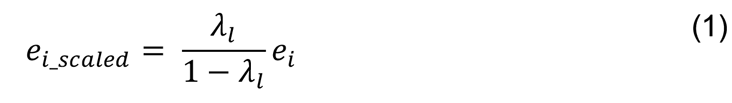

This scaling is equivalent to estimating diffusion distances from 1, 2. … ∞ steps. See section

“Measuring distances between cells in the phenotypic manifold” under the Palantir algorithm description for details on scaling and its impact on the representation. The number of components were chosen based on the Eigen gap of the Eigenvalue decomposition of the diffusion operator. The set of diffusion components is the same set used for running Palantir. Using diffusion components as inputs led to maps more representative of differentiation when compared to the maps generated on principal components or force directed graphs (Supp. Fig. 16). We found that force directed graphs represent the distinct mature populations better and provide less resolution in the regions of manifold where lineage decisions are being made. An example of generating tSNE maps using diffusion components is available here: http://nbviewer.jupyter.org/github/dpeerlab/Palantir/blob/master/notebooks/Palantir_sample_notebook.ipynb

#### Differential expression of genes

Differentially expressed genes between clusters were determined using MAST ^65^. MAST was run using default parameters with normalized counts (without log transform) as the input. Genes with FDR corrected p-value < 1e-2 and absolute log fold change > 1.25 were considered significantly different.

### The Palantir algorithm

#### Constructing a nearest neighbor graph representing the phenotypic manifold

Palantir first constructs a nearest-neighbor graph representing the phenotypic manifold, where each cell is connected to its most similar cells. Key to the success of this approach is that the resulting graph neighbors consist of cells in similar developmental states and that longer paths correspond to developmental trajectories. Given the extensive degree of sparsity and noise in scRNA-seq, finding nearest neighbors in the raw data using a simple similarity metric is likely to accumulate spurious connections and obscure the structure we are seeking.

To construct the neighbor graph based on robust trends in the data, Palantir uses diffusion maps ^21^, which project the data onto a low dimensional manifold that approximates the differentiation landscape. Diffusion maps have been previously used to study differentiation in single cell data ^2, 3^ and are particularly adept at capturing differentiation. Diffusion maps generate a low-dimensional embedding by approximating all possible paths via random walks through the graph, which effectively capture the major axes of variation in the data (Supp. Fig. 1).

The first step in constructing diffusion maps is to define a measure of similarity between cells. Following ^66^, we use an adaptive (width) Gaussian kernel to convert distances into affinities, so that similarity between two cells decreases exponentially with their distance. Typically, an isotropic or non-adaptive Gaussian kernel is used to measure the similarity with an inherent assumption the density of the data is uniform along the trajectory. However previous single cell studies have shown that while differentiation trajectories are continuous, they are punctuated by large changes in densities ^1, 2^ possibly representing meta-stable states. A non-adaptive kernel would be strongly biased by the densest regions. The adaptive kernel ^66^corrects for the densities by using the distance to the *l^th^* nearest neighbor as a scaling factor, thus equalizing the effective number of neighbors for each cell.

Formally, given a dataset, ***X*** ∊ ***R**^N×M^*, with *N* cells and *M* genes, a *k*-nearest neighbor graph, 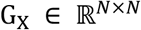 is constructed using the Euclidean distance. The distances are converted to affinities using the adaptive kernel as defined below.

The scaling factor of cell *i* is determined by

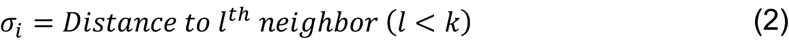

Given this, the similarity measure between two cells *i* and *j* is given by

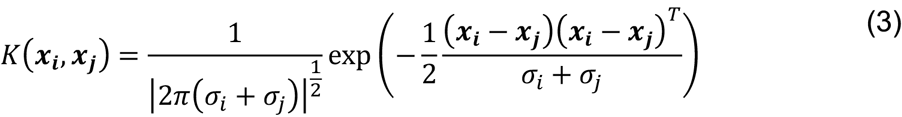

Where *x_i_* is the vector of gene expression for cell *i*. Thus, the above adaptive anisotropic kernel is used to define an affinity matrix, 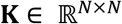 from the data. We then compute the Laplacian of the affinity matrix ***K*** to derive the diffusion operator 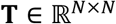, where *T_ij_* represents the probability of reaching cell *j* from cell *i* in one step. The Eigenvectors of the diffusion operator ***T*** are termed diffusion components and these represent major axes of variation in the underlying manifold from which the data was sampled. The top diffusion components (Eigenvectors of T) define a non-linear low dimensional embedding that approximates the phenotypic manifold of the data (Supp. Fig. 2) ^21^.

#### Pseudo-time ordering of cells

Once the manifold is constructed, the next step is to infer a pseudo-time for all cells in the data. The computed pseudo-time does not represent a single trajectory, but rather assigns each cell their relative distance from a starting cell, regardless of their lineage or terminal fates. Typically, diffusion maps have been used to characterize pseudo-time ordering of cells, constructing separate trajectories for each major axes of variation, based on individual diffusion components (DCs) ^3, 23, 24^. While a single DC can sometimes offer a reasonable approximation of an ordering towards a specific fate, we often observe many to many relationships between DCs and ordering leading to each terminal fate (Supp. Fig. 2). Therefore, Palantir takes multiple DCs into account when computing the pseudo-time of cells.

The embedded space (diffusion map) is used as an approximation of the differentiation landscape. Palantir uses Euclidean distance at multiple scales or multi-scale distance (elaborated below) in this embedded space to construct a more reliable nearest-neighbor graph, 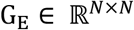 that filters out much of the noise in the original neighbor graph G_X_ (Supp. Fig 1a). Then pseudo-time is determined using shortest path distances in the graph G_E_ (Supp. Fig. 1b), since shortest path lengths better approximate the geodesic distances in the manifold ^67^.

The extremes of the diffusion components determine the boundaries of the phenotypic space and the start cell is defined as the boundary cell closest to the user defined starting point. Then, pseudo-time is initialized as the shortest path distances from this start cell. A shortcoming of shortest path distance is that it tends to accumulate noise with increasing distances ^1, 2^, thus, similarly to ^1, 2^, waypoints are used to refine the pseudo-time, defining the ordering based on a weighted vote of waypoints. Waypoints act as guides: the waypoint closest to the cell gets the highest vote in determining the position of the cell along pseudo-time. Thus, the success of Palantir requires that all regions of the manifold are well covered by waypoints that can guide the positioning of cells in their respective regions.

Our previous pseudo-time algorithms ^1, 2^ used a random sample of cells as waypoints. However, random sampling does not successfully cover the landscape in complex datasets, with multiple branches and the variable densities of cells along different lineages. Therefore, Palantir uses max-min sampling ^68^, an iterative procedure to choose waypoints that spread over and represent the entire manifold, rather than representing only the regions of high density.

In summary, the positioning is initialized based on shortest path distances from the start cell and is iteratively refined using the waypoints to fine tune the distances of the cells within the region of each waypoint. A weighted average across all waypoints is used to ensure the computation of a consistent global structure. Convergence of this procedure defines a final pseudo-time ordering of cells (Supp. Fig. 1c). Below we provide more detail:

#### Measuring distances between cells using multi-scale distance

Let the manifold be represented by 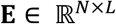 where *L* is the dimension of the embedding with *L* < *M*. The dimension *L* of the embedding is chosen using an Eigen gap among the top Eigenvectors. Let *λ*_1_,*λ*_2_…*λ_L_* be the corresponding Eigenvalues associated with diffusion components that define the manifold.

Given this, the distance between cells *i* and, *j* known as the diffusion distance, is defined by

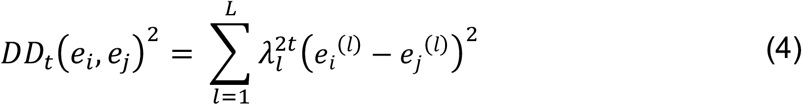

where *t* is the number of steps through the graph and *e_i(l)_* is the embedding of cell *i* along diffusion component *l*. Different stages of differentiation happen at different rates and occur at different densities in the population, thus a single *t* is unsuitable across the entire population. To avoid setting a particular *t*, in a similar manner to ^3^, we use multi-scale distance that accounts for all scales:

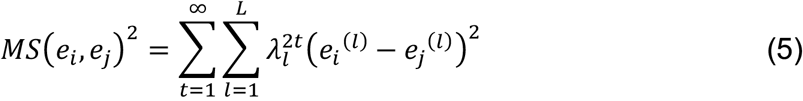

By definition, 1 > *λ*_1_ > *λ*_2_>…> *λ_L_* > 0, thus Equation (5) can be rewritten as

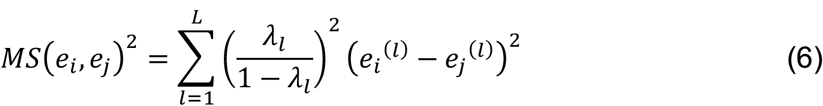

The use of multi-scale distance avoids the selection of an additional parameter (*t*) and also renders the distance robust to different choices of *L* (Supp. Fig. 17a-b), robust to outlier cells and density differences.

#### Max-min Waypoint sampling

Max-min sampling is an iterative procedure, where at each iteration, the chosen waypoint maximizes the minimum distance to the set of current waypoints ^68^, thus covering a new region of the manifold. Palantir uses max-min sampling along each diffusion component to sample waypoints.

Let **E**^(l)^ be the *l^th^* diffusion component. Max-min sampling is initialized with a randomly sampled cell from the diffusion component: *WS*^(l)^ = *Random*(*N*, 1). Distances along the component to the current waypoint set are computed for all the cells
initialized as the shortest path distances from

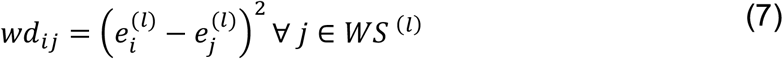

For each cell *i*, minimum of the current waypoint distances is computed

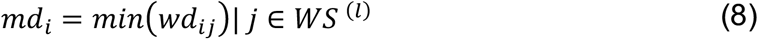

The cell with the maximum of these minimum distances is added to the waypoint set

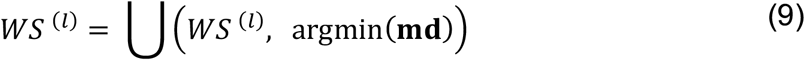

This procedure is repeated until the desired number of waypoints is sampled along the component and then repeated for all components. Union of the waypoints sampled along all diffusion components represents the final waypoint set, *WS*. An example of waypoint sampling along a component is shown in Supp. Fig. 17c.

#### Iterative pseudo-time computation

Palantir begins with designating a start cell based on a user defined starting point. It is assumed that the starting cell would reside at the boundary of the manifold, that is a cell that projects onto an extreme endpoint along of one of the diffusion components. First, the set of boundary cells is determined using

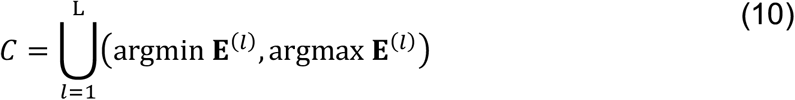

The extreme cell closest to the user input early cell *s* is then used as the start of the pseudotime, *s′*.

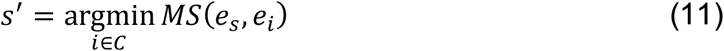

The pseudo-time, τ^(0)^, is initialized as the shortest path distances from the start cell *s*′. Shortest path distances are computed from each of the waypoints to all cells (Supp. Fig. 17d). These distances are then aligned to the start cell distances to compute waypoint perspectives (Supp. Fig. 17e). The pseudo-time is then updated as the weighted average of the different waypoint perspectives, ensuring that the pseudo-time of a cell is most strongly influenced by the waypoints closest to it, while maintaining a consistent global structure.

Formally, let *D_wi_* be the shortest path distance of cell *i* from to waypoint *w*. The perspective of a cell *i* relative to waypoint *w* is the distance of from early cell *s′*. is computed as

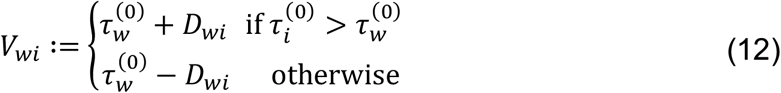

Note that the perspective of the early cell *s′* is the initial ordering τ^(0)^ itself.

The weighted average of waypoint perspectives is used to refine the pseudo-time, using an exponential weighting scheme where the weight is inversely proportional to the distance between the waypoint and the cell. The weights are determined as follows

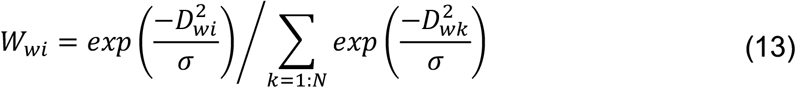

where *σ* is the standard deviation of distance matrix **D**. This defines the weight matrix 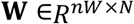. The weighted average is then calculated by

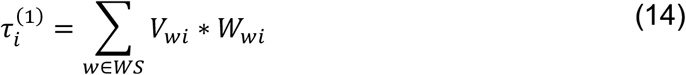

Note that the waypoints themselves are also cells and thus their relative distance to the start cell is modified and updated by this procedure. The updated ordering is then iteratively refined until convergence to obtain a final pseudo-time, *τ* (Supp. Fig. 1c).

### Inferring the Terminal Fates and differentiation potential

#### Modeling Differentiation as an Absorbing Markov Chain

Consider the neighbor-graph spanning the waypoints, G′_E_ ∊ G_E_. Differentiation is modeled as a stochastic process, implemented as a Markov chain, where a cell reaches one or more terminal states through a series of steps in the manifold (Fig. 1b), based on the assumption that paths in the neighbor-graph G_E_ correspond to possible differentiation paths. However, differentiation is a directed process, from a less differentiated to a more differentiated state, whereas G′_E_ is an undirected graph.

The inferred pseudo-time τ provides directionality that can be used to orient neighboring edges in G′_E_, thus allowing construction of a directed graph for the Markov chain. A naïve approach would prune all edges that violate the pseudo-time order to prevent de-differentiation paths. However, there is uncertainty in the pseudo-time estimate of the cells and therefore we use the estimated scaling factor for each cell in Equation 2 as a measure of the uncertainty in the pseudo-time estimate. Specifically, an undirected edge between cell *i* and its neighbor cell *j* is converted to a directed edge from cell *i* to cell *j* if *τ_i_* < *τ_j_*. The edge between cell *i* to cell *j* is pruned if *τ_i_* > *τ_j_* and the distance between the two cells exceeds the scaling factor of cell *i* determined using Equation 2 (Supp. Fig. 1d-h).

Formally, undirected graph in the manifold, G_E_ is converted to directed graph, 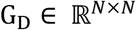 using

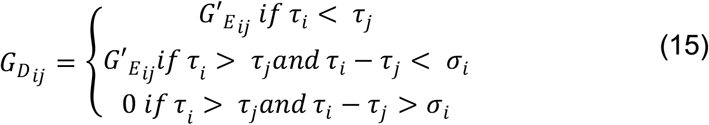

These distances are then converted to transition probabilities to construct the Markov chain. First, distances are transformed to an affinity matrix 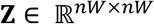 using the kernel defined in Equation (3) where *nW* is the number of waypoints. These affinities can be converted to probability matrix by dividing each affinity by the degree of the node in Z representing that cell.

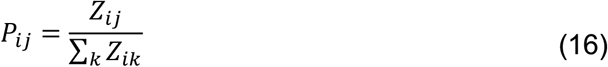

The transition probability matrix **P** represents the Markov chain of the manifold, where *P_ij_* represents the probability of reaching a cell in state *j* from a cell in state *i* in one step. As a first degree of approximation, our approach assumes that this probability of transition corresponds to the degree of cell state similarity between *i* and *j*. While development is a closely regulated process, at these very close distances, stochastic molecular processes of degradation and transcription likely play a significant role in. At longer distances, the regulatory processes driving development are implicitly encoded in the defined structure of the manifold graph G_D_. That is, the probability of reaching a cell in state *j* from *a more distinct cell* in state *i* is computed over the course of *many steps* and will be high if *many paths* connect them, i.e., there is high density of *observed* intermediary cell states between them.

By definition terminal states are not expected to differentiate further, thus to ensure that the random walks terminate when a terminal state is reached, all outgoing edges are removed from terminal states. Terminal states can be externally defined based on prior knowledge or can be computationally derived directly from the Markov chain using no additional knowledge, as we describe below. Given a set of terminal states *TS*, we convert the Markov chain P into an absorbing Markov chain A by setting the terminal states as absorbing states i.e., a state with no outgoing edges.

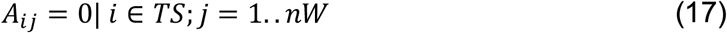

#### Identifying Terminal States

The graph structure and its associated Markov chain can be used to infer the terminal states directly from the data, using only the initial starting point as prior information. In the Markov chain P, random walks tend to move in the direction of the terminal states. Since a pseudo-time underlies the Markov chain, we expect random walks to converge into the terminal states at the boundaries of the manifold. If the graph construction were perfect, we expect that these terminal states have no outgoing edges and thus be absorbing states. However, the chain was constructed with implicit uncertainty (e.g. the backward edges within the range of the scaling factor) and is therefore imperfect. Nevertheless, as the random walks are directed towards the terminal states, the steady state distribution of the Markov chain is expected to impart high probabilities to terminal states and states proximal to them as opposed to the intermediate states. Thus, Palantir identifies terminal states as extrema of diffusion components (boundary cells, *C*), that are also outliers in the steady state distribution of the Markov chain (Supp. Fig. 18).

The stationary distribution is the probability distribution over the states of the Markov chain that remains invariant as time progresses, i.e. the steady state distribution. Formally, if **π** represents the stationary distribution, then **π** = **P** * **π**. The first *left* Eigen vector of the Markov chain P represents the stationary distribution and is thus easy to compute. The outliers in this distribution can be identified using the Gaussian percent point function (i.e., inverse of the cumulative distribution function) using the median absolute deviation of the stationary distribution as the scale. Median absolute deviation is a robust measure of variance in univariate data ^69^. Let π represent the stationary distribution. The median absolute deviation is computed as

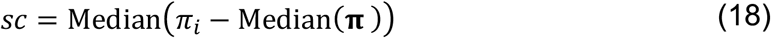

The outliers are identified as

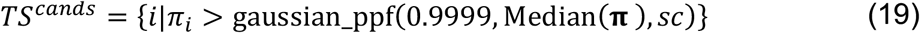

This threshold robustly identifies the different terminal states across the different data sets. The set of states in *TS^cands^* that are also diffusion component extremes are chosen as the terminal states of the system (Supp. Fig. 18, Fig 1c).

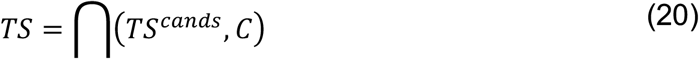

#### Cell fate/differentiation potential characterization

Random walks through the Markov chain between intermediate and terminal states can be used to compute the probability of a cell starting at an intermediate state reaching the corresponding terminal state. For each cell, we wish to calculate its branch probability vector **B**_*i*_, denoting the probabilities it might reach each of *b* absorbing terminal states. An advantage in modeling differentiation as an absorbing Markov chain is that the branch probabilities can be computed as follows:

The absorbing Markov chain **A** can be represented as

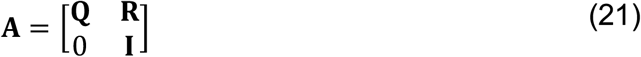

Where **Q** is a (*nW* – *b*) × (*nW* – *b*) matrix of transition probabilities between intermediate states, **R** is a (*nW* – *b*) × *b* matrix of probabilities between intermediate states and terminal states and I is a *b* × *b* identity matrix.

Next the fundamental matrix **F** is computed using

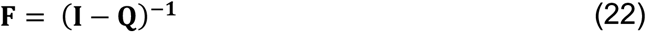

*F_ij_* represents the probability of reaching intermediate state *j* from another intermediate state *i* in *1,2,…∞* steps

The fundamental matrix is then used to compute the differentiation probabilities

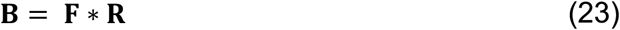

where *B_ij_* represents the probability of cell in intermediate state *i* reaching the terminal state *j* in *1,2,…∞* steps. **B***_i_* is a multinomial probability distribution such that ∑*_j_ B_ij_* = 1. The branch probabilities of terminal states are set as follows.

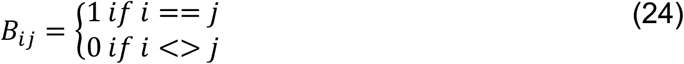

The waypoint branch probabilities are projected onto all the cells using weighting scheme defined in Equation (14)

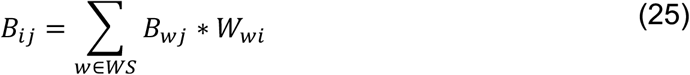

Finally, we define the differentiation potential of each state to be the entropy of the branch probability vector **B***_i_* and this captures the degree of uncertainty in final terminal state (Fig. 1d).

Differentiation potential is a quantitative measure of the cell fate plasticity and represents the potential set of terminal states that a cell in an intermediate state can reach. Greater the entropy, higher is the number of terminal states the can potentially be reached by the cell in a particular state. As a result, the cells at the beginning of the pseudo-time are associated with the highest differentiation potential (entropy) (Fig. d(1)) whereas cells close to terminal states have the lowest differentiation potential (Fig. d(3, 7)). Crucially, differentiation potential captures the continuity in cell fate determination (Fig. 1d(2, 4-6)) and is a better representation of the differentiation processes as opposed to well-defined branch points. In summary, Palantir characterizes the continuity in both cell state and cell fate by modeling differentiation as a stochastic process.

#### Gene expression trends along lineages

Palantir’s pseudotime represents an ordering over all cells across all lineages and provides the position of all the cells relative to the start cell. In addition, Palantir branch probabilities represent the probability of a cell, in any state, to reaching each of the terminal states. Therefore, Palantir’s ordering and branch probabilities represent a unified framework that enables computation and comparison of gene expression trends across the different lineages. This framework is used to compute gene expression trends for each lineage as follows: rather than segmenting the cells that belong to each lineage, the trend is computed using *all* the cells, each weighted by its probability to belong to that particular lineage. Cells that are not committed to a particular lineage can provide input to multiple lineages, whereas low probabilities naturally exclude cells that belong to unrelated lineages.

We take two approaches to improve the robustness and resolution of the computed trends: MAGIC ^66^ to impute missing values and generalized additive models ^25^ to determine robust trends. Gene expression trends are computed using MAGIC imputed ^66^ data to prevent dropouts from adversely affecting the trends. MAGIC imputes missing values for each cell based on cells that are most similar to it by using the covariate relationships between genes. Sliding window approaches on the other hand, average expression over many cells in a univariate manner, regardless of other genes. MAGIC, like Palantir is also based on diffusion maps and we use the same diffusion operator for both MAGIC and Palantir. We note that imputed data is only used to compute the expression trends after Palantir pseudo-time and branch probabilities have been computed using the non-imputed data.

Sliding window approaches are sensitive to density differences even with imputed data (Supp. Fig. 15d). We therefore used generalized additive models (GAMs) to determine gene expression trends along each lineage (Supp. Fig. 3), increasing robustness and rendering trends less sensitive to changes in cell density along the lineage. The gene expression trend for gene *g* and branch *b* is fit using

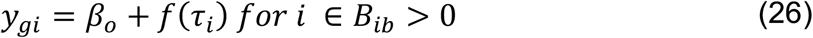

where *y_gi_* is the expression of gene *g* in cell *i* and *τ_i_* is the pseudotime ordering of cell *i*. Cubic splines are used as the smoothing functions since they are effective in capturing non-linear relationships ^25^.

The pseudotime is then divided into 500 equally sized bins and the smooth trend is derived by using the fit from Equation 25 to predict the expression of the gene at each bin (Supp. Fig. 3). The standard deviations of expression along each bin is determined by the standard deviation of the residuals of the fit and is computed as follows

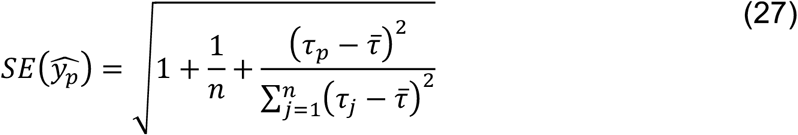

where 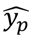 is the predicted expression at bin *p* and *τ̄* is the average pseudo-time across all cells. The computation and plotting of gene expression trends are demonstrated here: http://nbviewer.jupyter.org/github/dpeerlab/Palantir/blob/master/notebooks/Palantir_sample_notebook.ipynb

### Single cell transcription factor activities

#### Bulk ATAC-seq data processing

Bulk ATAC-seq data was downloaded from GEO (GSE75384) Reads were aligned to hg38 genome using bowtie2. PCR duplicates were removed using samtools, rmdup. Reads with fragment size < 150 bp, representing TF binding events ^70^ were retained for downstream analysis. After size selection, reads were pooled from all cell types and replicates. Only the first read from the pair was used for peak calling since a single transposase nick is sufficient proof for exposed chromatin ^70^. Peak calling was performed using macs2 with a permissive p-value threshold of 1e-5 and with the parameter “nomodel” turned on to prevent shifting of positive and negative reads towards each other ^47^. IDR ^71^ was then used to identify reproducible peaks for each cell type: IDR was performed on each pair of available replicates and a peak was assigned to a cell type if the peak passed IDR < 0.1 in at least 50% of the replicate comparisons.

#### Motif discovery

SeqGL ^47^ was run separately for each cell type with default parameters using the reproducible peaks for the respective cell type. SeqGL outputs a predicted sequence affinity for each TF, peak pair. The sequence affinity represents a quantitative measure of the k-mer sequence preferences: a higher value represents a greater chance that the TF binds at the genomic location spanned by the ATAC-seq peak.

#### Single cell TF activity

ATAC-seq peaks were assigned to the gene with the nearest transcription start site, which is a reasonable approximation of enhancer target assignment in the absence of chromosome interaction data ^11^. Sequence affinities for all TF-gene pairs were determined by aggregating the affinities across all peaks assigned to gene. Recent studies have shown that these affinities correlate strongly with expression change of the targets indicating that the sequence affinities approximate the regulatory effect of a TF on its target ^11^. Therefore, correlation between target expression and predicted TF sequence affinity was used as the TF activity for each cell (Fig. 4b). The activities were determined separately for promoter peaks (peaks within 2kb of the transcription start site) and enhancer peaks (peaks at distance > 2kb of the transcription start site). As a demonstration of the importance of cell type context for determining TF targets, Supp. Fig. 12h shows TF activity trends for Runx in different cell types. The targets of Runx, a transcriptional activator, show higher expression in the corresponding cell type, demonstrating the accuracy of computing TF activities using correlation between target expression and predicted sequence affinities

### Subsampled data used for figure 1

A dataset was generated using the human CD34+ hematopoiesis dataset by waypoint sampling of cells from erythroid and myeloid lineages (clusters 0, 1, 2, 3, 4, 6, 7, 8 - Supp. Fig. 4a). tSNE map was generated as described in “Single cell RNA-seq data preprocessing” and the projection of stem cells was manually adjusted for cleaner visualization.

### Application of Palantir to CD34+ cells

Palantir was applied to each replicate separately using 1200 waypoints and one of the CD34+ cells as the start cell. The parameter *k* was set to 10% of the total number of cells in the data. The results however are stable to the choice of *k* (Supp. Fig. 5). The number of diffusion components were chosen based on the Eigen gap of the Eigenvector decomposition of the diffusion operator. The results are stable to choice of the number of diffusion components and the choice of waypoints (Supp. Fig. 5).

#### Robustness of Palantir results to parameters

Palantir has the following parameters or variables: (a) *k*, number of neighbors for nearest neighbor graph, (b) Waypoint samples and (c) Number of diffusion components, which by default is determined based on the Eigen gap. Palantir’s robustness to these parameters was tested using data from replicate 1 of the CD34+ bone marrow data. Palantir was run with different parameters and the robustness of the results was measured using Pearson correlation of both pseudo-time and differentiation potential between a given pair of Palantir runs. The same start cell was used as input for each run.

Robustness to *k* was tested by fixing the number of diffusion components, waypoints and terminal states. Robustness to number of diffusion components was tested by using fixing *k*, waypoints and terminal states. Robustness to waypoint sampling was tested by fixing *k* and the number of diffusion components. Palantir results are very robust to all the different parameter settings (Supp. Fig. 5).

#### Comparison of Palantir results across replicates

Palantir results, specifically pseudo-time and differentiation potential, from one replicate are projected onto cells from a second replicate using mutually nearest neighbors (Supp. Fig. 8). The projected results are then correlated with Palantir results derived *de novo* from the second replicate to measure reproducibility of Palantir results across the replicates.

Let *N_1_* and *N_2_* be the number of the cells in replicate *1* and *2* respectively. As a first step, the count matrices of both replicates are combined to create a unified molecule count matrix using genes detected in both replicates. This matrix is normalized as described in “Single cell RNA-seq analysis: Data preprocessing” and log transformed with a pseudo count of 0.1, followed by PCA. Principal component space of the combined count matrix is used to determine the k-nearest replicate *1* neighbors of replicate *2*. This neighborhood graph can be represented by an adjacency matrix *D*^21^ ∊ *R*^*N*_2_×*N*_1_^ where 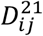 is the distance between cell *i* of replicate *2* and cell *j* of replicate *1* if *i* and *j* are neighbors. Similarly let *D*^12^ ∊ *R*^*N*_1_×*N*_2_^ represent the adjacency matrix of replicate *2* neighbors of replicate *1*.

Mutually nearest neighbors between the two replicates is computed as below

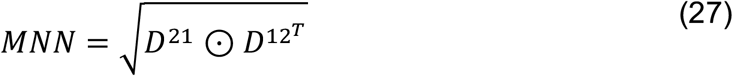

where *MNN* ∊ *R*^*N*_1_×*N*_2_^ and ʘ is the Hadamard product or element-wise multiplication operator.

The distances of the *MNN* adjacency matrix is converted to an affinity matrix using Equation 13.

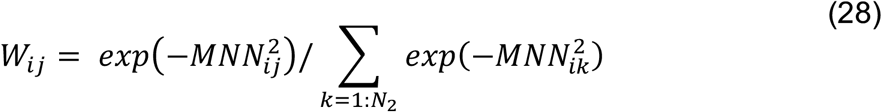

Palantir results of replicate *1* are projected on to the cells of replicate *2* using the weights computed in Equation 28. The projected results are thus a weighted average of the mutually nearest neighbors of each cell.

Let *τ*_*Rep*1_ and *τ*_*Rep*2_ be the *de novo* pseudotime ordering of replicates 1 and 2 respectively. The projected pseudo-time is computed as follows

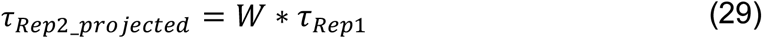

Pearson correlation between *τ*_*Rep*2_*prjected*_ and *τ*_*Rep*2_ gives a measure of reproducibility of Palantir pseudo-time. Similarly, the projected differentiation potential is computed as follow

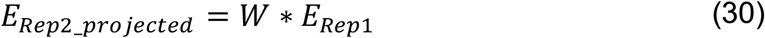

Similar to the pseudo-time, Pearson correlation between *E*_*Rep*2_*prjected*_and *E*_*Rep*2_ gives a measure of reproducibility of the differentiation potential.

#### Clustering of gene expression trends

Genes were selected based on significant differential expression as determined by MAST (FDR corrected p-value < 1e-2 and absolute log fold change > 1.25). Genes that were significantly high or low in stem and precursor cell clusters (0 or 1: 2176 genes) (Supp. Fig. 4a) were used for analysis in Fig. 3b and genes that were significantly high or low in early cell and erythroid cell clusters (0, 1, 2 or 8: 3322 genes) (Supp. Fig. 4a) were used analysis in Fig. 3c-d. Gene expression trends were z-transformed to put them on the same scale and clustered using Phenograph ^20^ (Supp. Fig. 10). A high-value of *k* (150) was used to avoid over-fragmentation of the gene trend clusters. The within cluster sum-of-squares each trend cluster was significantly lower for clusters derived from Phenograph when compared to trend matching techniques such as dynamic time warping. Gene ontology analysis was performed to annotate each cluster, measuring enrichment using the hypergeometric test. The following gene sets from Molecular Signature Database (MSigDB) (http://software.broadinstitute.org/gsea/msigdb/index.jsp) were tested: (a) c5 GO biological process gene set, (b) H hallmark gene sets and (c) c2 canonical pathway gene sets.

In order to compare the change of differentiation potential relative to gene expression changes, cells were divided into equal sized bins along the Palantir pseudotime ordering. Mean expression of genes from the relevant clusters were determined to generate the histograms in Fig 3. The point of maximal differentiation potential change was determined using the second derivative. Stem and precursor cells (clusters 0 and 1) were used for analysis in Fig. 3b whereas these cells along the erythroid lineage (clusters 0, 2 and 8) were used for analysis in Fig. 3c-e. Similar analysis was performed for replicates 2 and 3 using the genes that were differential in replicate 1 (Supp. Fig. 11).

The computation and plotting of gene expression trend clustering for a particular lineage is shown here: http://nbviewer.jupyter.org/github/dpeerlab/Palantir/blob/master/notebooks/Palantir_sample_notebook.ipynb

#### GATA2 identification

We downloaded the list of human TFs from AnimalTFDb ^72^. TFs that were significantly high in one of the erythroid clusters (clusters 2 or 8 - Supp. Fig. 4a) were annotated as erythroid TFs. We reason that a TF that potentially plays a role in lineage specification should correlate with the branch probability during the specification phase i.e., when the branch probability begins to increase along pseudo-time. We used the second derivative of the erythroid probability trend to approximate the point along the pseudo-time where there is a switch from lineage specification to functional commitment, since the second derivative indicates the point of maximal change in the trend (Supp. Fig. 12b).

We computed the correlation between TF expression trend and the erythroid probability trend for each erythroid TF defined above (Supp. Fig. 12b). To avoid down-weighting TFs that are potentially downregulated following commitment, during the functional specification phase, we computed the correlation until the point of maximal differentiation potential change (and not along the entire pseudo-time). Only TFs with both a high correlation and with sufficiently high expression levels in early cells were considered candidate erythroid specifiers. The comparison of branch probability correlation and mean expression in stem cell cluster (cluster 0 - Supp. Fig. 4a) shows that GATA2 is a clear outlier (Supp. Fig. 12b). GATA2 is also the only factor with high correlation and high progenitor cell expression for which a motif was identified in bulk ATAC-seq data.

### Additional datasets

#### Mouse hematopoiesis dataset

Mouse hematopoiesis dataset ^6^ was downloaded and preprocessed using the procedure outlined in scanpy (https://github.com/theislab/paga/blob/master/blood/paul15/paul15.ipynb). A cluster of cells annotated as DCs were projected as a clear outlier along a diffusion component without a well-defined differentiation path (likely due to insufficient cell sampling) and therefore were excluded from the analysis. PCA was performed on the preprocessed data and components that explain 85% of the variance were used for generating diffusion maps as described in “The Palantir algorithm”. Eigen gap suggested use of 7 diffusion components, but 13 components were used instead to ensure inclusion of all cell types. Note that the frequencies of some of the populations such as basophils is extremely low necessitating the inclusion of additional components.

Palantir was run using one of the cells annotated as MEPs since these are the most primitive cells present in the data. Palantir automatically determined the different terminal states and determined pseudotime ordering, differentiation potential and branch probabilities. Differentiation potential trends and gene expression trends were generated as described in “Gene expression trends” section.

#### Mouse colon data

Raw counts for the mouse colon dataset ^48^ was downloaded from GEO (GSE102698 - https://www.ncbi.nlm.nih.gov/geo/query/acc.cgi?acc=GSE102698). Cells with low molecule count (<1000) and high mitochondrial molecule fraction (>0.2) were excluded from the analysis. Immune cells were also excluded since they are not relevant for differentiation. Data was normalized as described in “single cell data preprocessing”. Phenograph clustering of data revealed a cluster of cells with low molecule count distribution, which was excluded from the analysis. To maintain consistency with the analysis in the original publication, the data was not log transformed and was restricted to genes used by the authors. The gene list was downloaded from Flowrepository (http://flowrepository.org/id/FR-FCM-ZYAG).

As before, PCA was performed to reduce the data to 20 components (explaining 85% of the variance) and diffusion maps were computed using PCs as the input. Palantir was run using one of the Lgr5+ stem cells as the start. Palantir automatically identified colonocytes, goblet cells and Reg4+ goblet cells as the terminal states but failed to identify Tuft cells as one of the terminal states. Tuft cells are very similar in their expression profiles to the early cells and thus there was not sufficient variability for the small number of Tuft cells to be projected onto a distinct diffusion component (note, we believe greater cell numbers would have resolved this). The results in Fig. 5b were generated by manually setting Tuft cells as one of the terminal states.

### Performance of competing methods on the CD34+ marrow data

Supp. Fig. 14 details the performance of popular pseudotime methods like Diffusion Maps, Slingshot, Graph Abstraction, Monocle 2 on the CD34+ marrow data.

#### Diffusion maps

Individual diffusion components which explain ordering along a particular lineage were identified using correlation of the projections along the component with Palantir pseudotime although typically this is performed by visual inspection. Gene expression trends were computed by a sliding a window spanning 1% of cells along the diffusion component to compute the mean and the standard deviation (Supp. Fig. 14a).

#### Monocle 2

Monocle 2 uses a reverse graph embedding which simultaneously learns a principal graph that approximates the low dimensional manifold and projection of cells onto this graph to reconstruct single cell trajectories ^9^.

Monocle 2 was run with default parameters for UMI counts specified in the Monocle vignette (Supp. Fig. 14b). Specifically, Monocle 2 was run to embed the data into two dimensions. The structure and the states identified by Monocle 2 do not reconcile with the results obtained using Palantir, Slingshot and PAGA, nor with published hematopoiesis literature. The expression of marker genes on the embedding did not clearly indicate clear lineages or expression trends and therefore the trends for different lineage were not presented for Monocle.

#### Partition based Graph Abstraction (PAGA)

PAGA aims to reconcile clustering and trajectory inference and is particularly adept to scaling to large number of cells ^7^. PAGA generates an abstracted graph representing the differentiation tree underlying the data. The gene expression trends are fit by computing a pseudo time ordering for each lineage separately using diffusion pseudo time and then a moving average along the resulting pseudotime of cells. PAGA was run using default preprocessing steps outlined in https://github.com/theislab/paga/blob/master/blood/paul15/paul15.ipynb with log transformation of the normalized data (Supp. Fig. 14c).

#### Slingshot

Slingshot takes as input a clustering and low dimensional embedding of the data ^8^. Slingshot first determines a minimum spanning tree through the clusters to identify the overall branch structure of the data. Slingshot then fits principle curves for each branch/ lineage and uses orthogonal projections against these principle curves to determine the pseudotime ordering. Finally, Slingshot uses GAMs with loess fits to determine gene expression trends.

Slingshot was run with the clusters obtained from Phenograph (Supp. Fig. 4a) and diffusion components (Supp. Fig. 2b) generated using the adaptive kernel for comparison (Supp. Fig. 14d) using default parameters.

#### Software availability

Palantir is available as a python module here: http://github.com/dpeerlab/Palantir/. A jupyter notebook detailing the workflow including data preprocessing, running Palantir along with a demonstration of various plots and visualizations is available at http://nbviewer.jupyter.org/github/dpeerlab/Palantir/blob/master/notebooks/Palantirsamplenotebook

## Acknowledgments

We thank Roshan Sharma for valuable conversations related to this manuscript, Caitlin Trasande for helping write the manuscript and Elham Azizi, Cassandra Burdziak and Kat Hadjantonakis for valuable comments about the manuscript. This study was supported by NIH grants NIH DP1-HD084071, NIH R01CA164729, Cancer Center Support Grant P30 CA008748 and Gerry Center for Metastasis and Tumor Ecosystems.

## Author Contributions

M.S and D.P. conceived the study, designed and developed Palantir, developed additional analysis methods, analyzed the data and wrote the manuscript. M.S implemented Palantir and all other analysis methods. V.K. and L.M. designed, optimized and executed all single cell RNA-seq experiments. J.L and D.P developed an early theory on application of Markov chains to single cell data. M.S and A.G developed trend based clustering analysis.

## Supplementary figures

**Supp. Fig.1:**
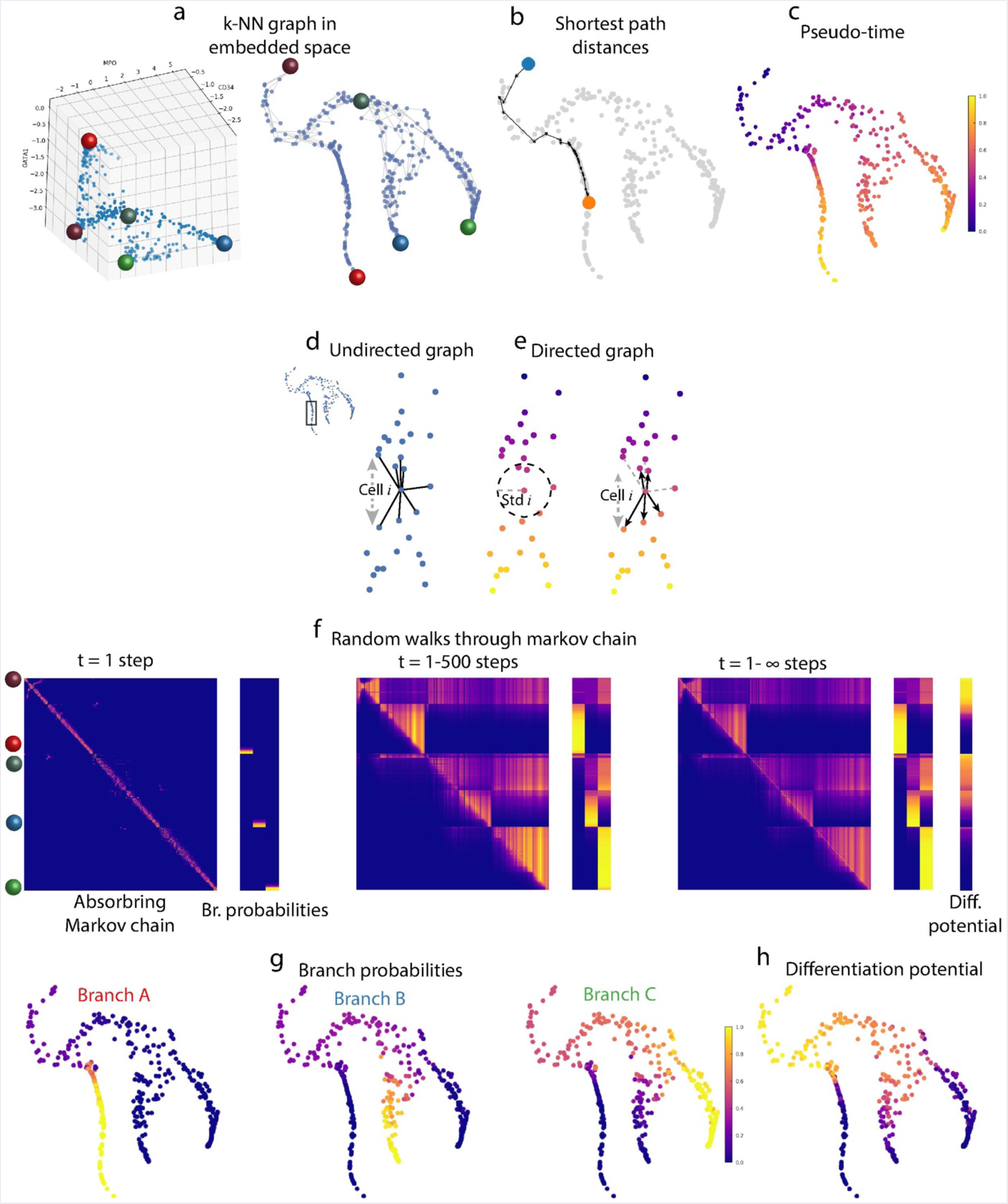
Palantir algorithm outline. Illustration of steps in the Palantir algorithm, using the same data as Fig 1b. (a) High dimensional representation of the data, each dot represents a cell plotted based on expression of CD34 (x-axis), MPO (y-axis) and GATA1 (z-axis) (left panel). Right panel: Same tSNE plot as Fig. 1b generated using the diffusion components of the cells in the left panel. (b) Illustration of shortest path from blue cell to the orange cell. (c) Cells colored by Palantir pseudo-time. (d-e) Illustration of Markov chain construction. (d) Edges in the undirected graph can take cells both forward and back along pseudo-time. (e) The scaling factor associated with each cell (Equation 2) can be used as measure of uncertainty in the pseudo-time estimate (left panel). Edges that go backward beyond the pseudo-time uncertainty are pruned and the retained edges are converted to directed edges (right panel). (f) Heatmaps showing the evolution of absorbing Markov chain and branch probabilities for random walks of different lengths. (Left panel: 1 step, middle panel: 1…500 steps and right panel: 1…∞ steps). For each panel, rows and columns in the Markov chain heatmap represent all non-terminal cells ordered by Palantir pseudo-time. The value (*i,j*) represents the probability of cell *i* reaching cell *j* in the specified number of steps. Rows in the branch probabilities heatmap represent non-terminal cells and the columns represent the terminal states. The value (*i,j*) represents the probability of cell in non-terminal state *i* reaching terminal state *j* in the specified number of steps. The position of the individual cells highlighted in 1a are shown on the left. (g - h) Cells colored by Palantir branch probabilities and differentiation potential.

**Supp. Fig. 2:**
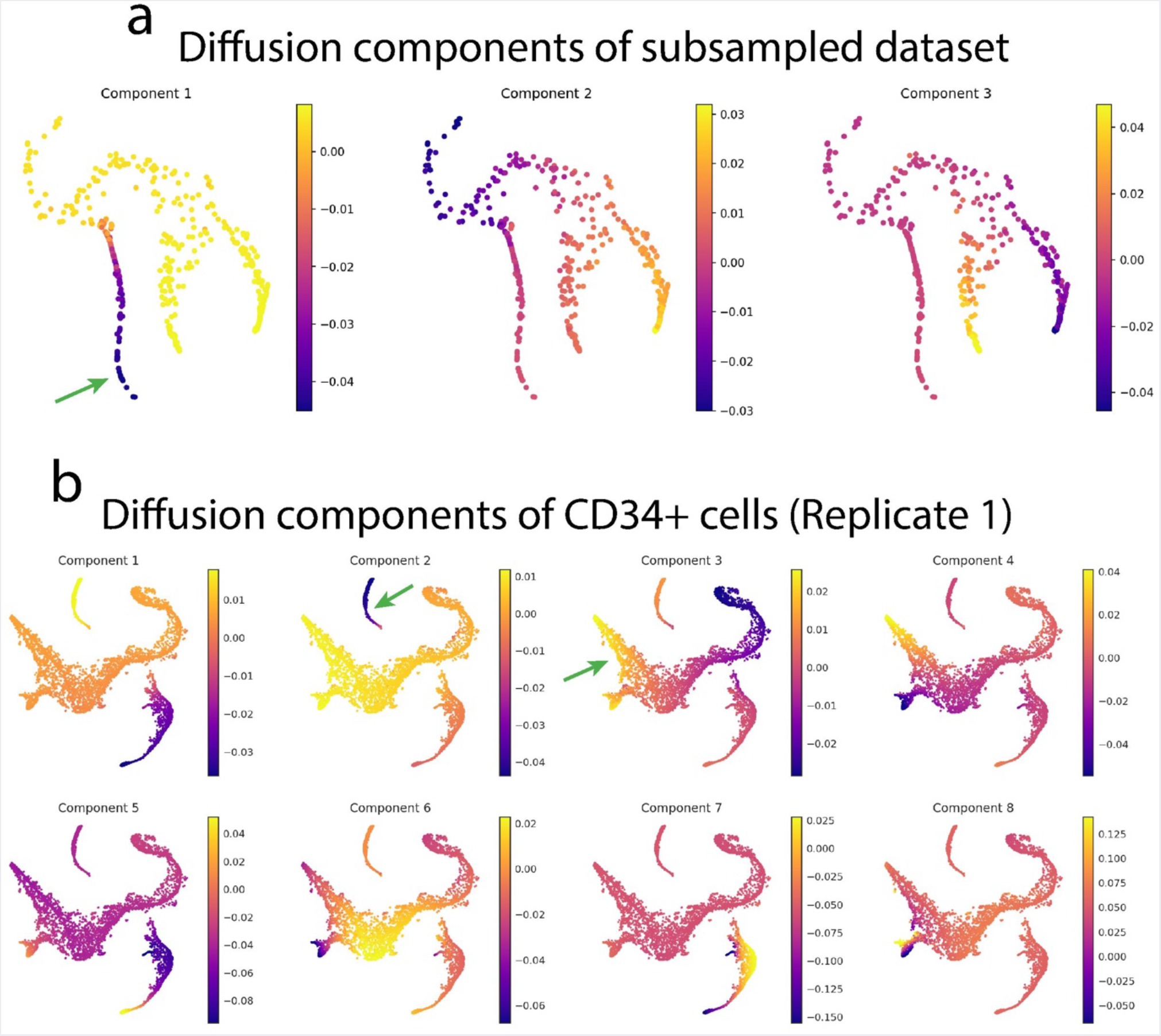
Diffusion components are not sufficient to represent pseudo-time for all lineages. (a) tSNE plots of the subsampled dataset used in Fig. 1 and Supp.Fig. 1, colored by diffusion components. (b) tSNE plots for CD34+ cells presented in figure 2, colored by diffusion components. Green arrows indicate the lineages for which ordering can be determined using a single component, whereas the ordering of the remaining lineages requires two or more components.

**Supp. Fig. 3:**
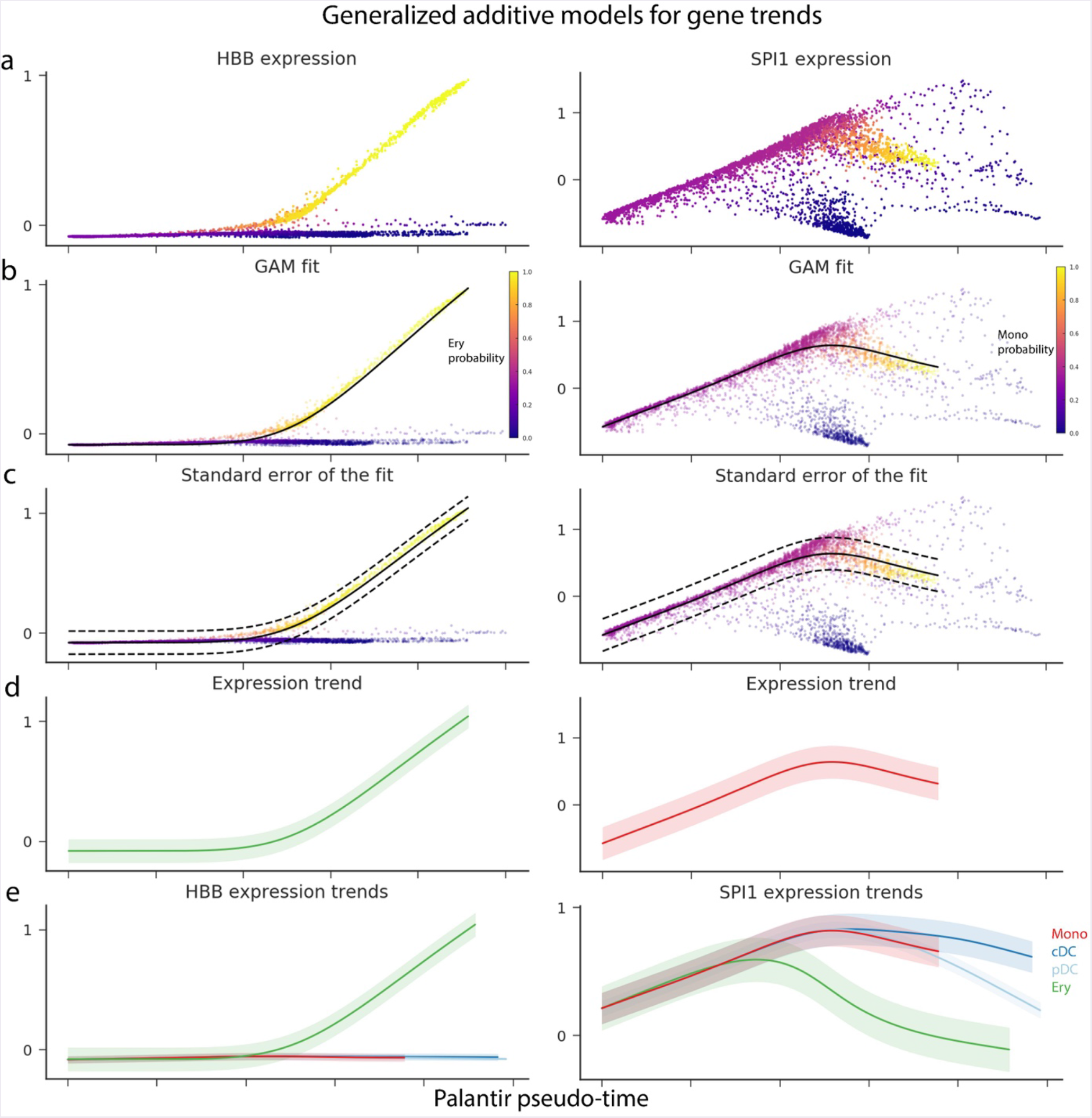
Characterizing gene expression dynamics. The characterization of gene expression dynamics is illustrated with two examples: HBB, a gene expressed specifically in the erythroid lineage (left panels) and SPI1, a gene with higher expression in the monocytic lineage (right panels) (a) Plots showing the MAGIC ^66^ imputed expression (y-axis) of HBB and SPI1 respectively along Palantir pseudo-time (x-axis). Cells are colored by the erythroid and monocyte branch probabilities respectively. (b) Same as (a) with the trend fit computed using Generalized Additive Models (GAMs) ^25^ shown in black. Each cell is weighted by the branch probability and thus no pre-selection of cells is necessary for computing trends along a particular lineage. (c) Same as (b) with standard deviations of the fit shown in dotted lines. (d) The expression trends are represented as a smooth fit with the standard deviation of the fit shown in a lighter shade. (e) Gene expression trends for HBB and SPI1 for the erythroid and myeloid lineages. For any gene, trends can be computed across all lineages since Palantir determines a single pseudotime across lineages.

**Supp. Fig. 4:**
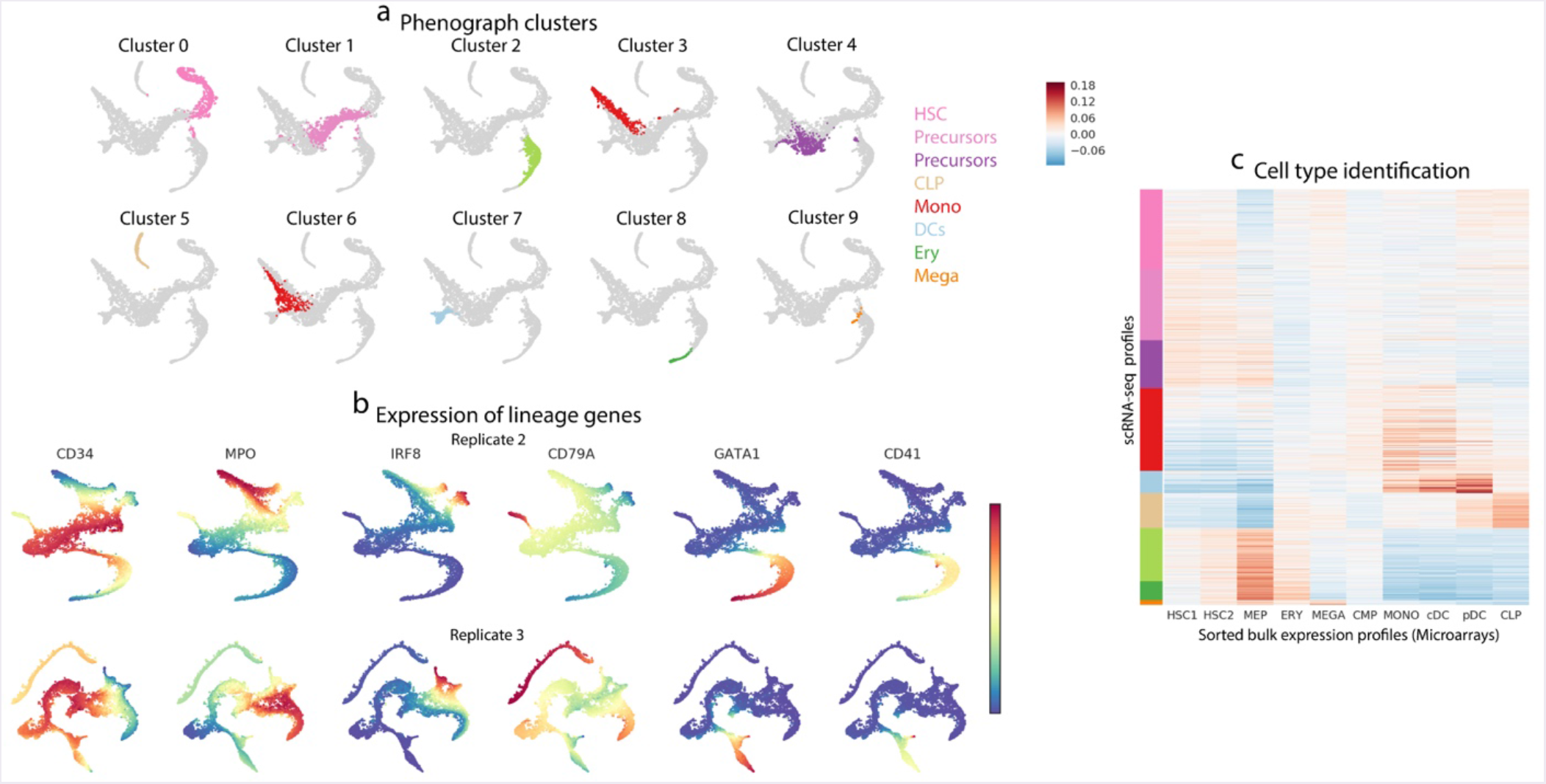
Cell lineages identified in CD34+ human bone marrow cells. (a) Replicate 1 CD34+ cells from human bone marrow colored by Phenograph clusters using the scheme presented in Fig. 2b. (b) Replicate 2 and 3 cells, colored by expression of lineage characteristic genes (Fig. 2f) demonstrating that the spectrum of lineages identified from CD34+ bone marrow cells is consistent across three independent human donors. (c) Heatmap of the correlation between bulk sorted expression profiles generated using microarrays and scRNA-seq profiles (Replicate 1). Rows represents single cell and columns represent bulk samples. Cells are ordered as in Fig. 2a. Median expression profiles from cell clusters (a) were correlated with bulk expression profiles to annotate clusters with cell types.

**Supp. Fig. 5:**
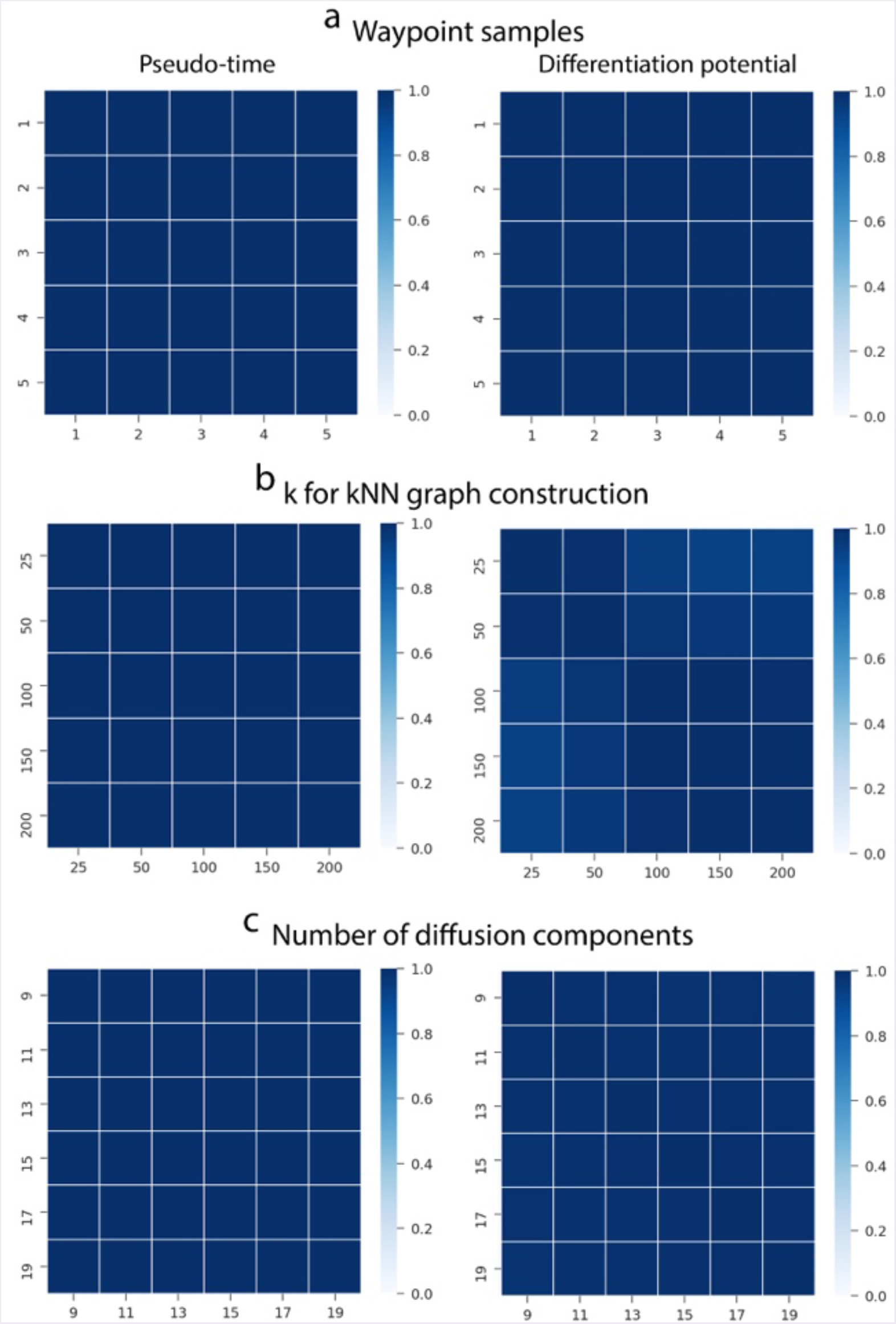
Palantir results are robust to different input parameters. Palantir robustness was measured by testing a range of different parameters using replicate 1 as the test bed. The results between two different runs were compared by determining the Pearson correlation between pseudo-time and differentiation potential. Each heatmap represents the correlation of either pseudo-time or DP between a pair of Palantir runs. Palantir results robust for (a) Different waypoint sampling with fixed number of waypoints, diffusion components and k for kNN graph construction (b) Different k for kNN graph construction with a fixed set of waypoints, diffusion components and terminal states (c) Different number of diffusion components with a fixed set of waypoints, k for kNN graph construction and terminal states

**Supp. Fig. 6:**
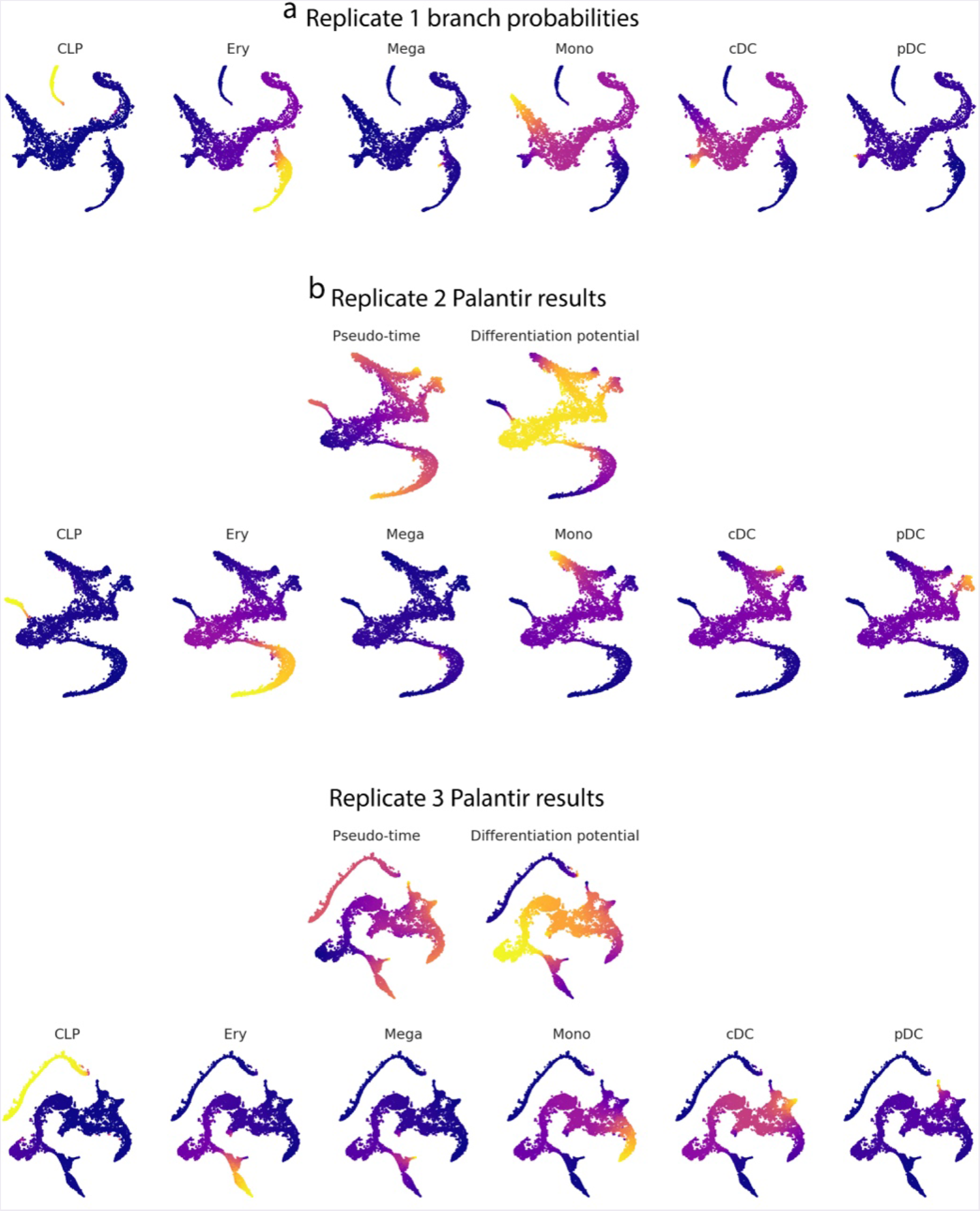
Palantir is reproducible across human bone marrow replicates. (a) Cells plotted on tSNE based on diffusion components and colored by Palantir branch probabilities for Replicate 1. (b-c) Cells plotted on tSNE based on diffusion components and colored by Palantir results: pseudo-time, differentiation potential and branch probabilities for replicates 2 and 3.

**Supp. Fig. 7:**
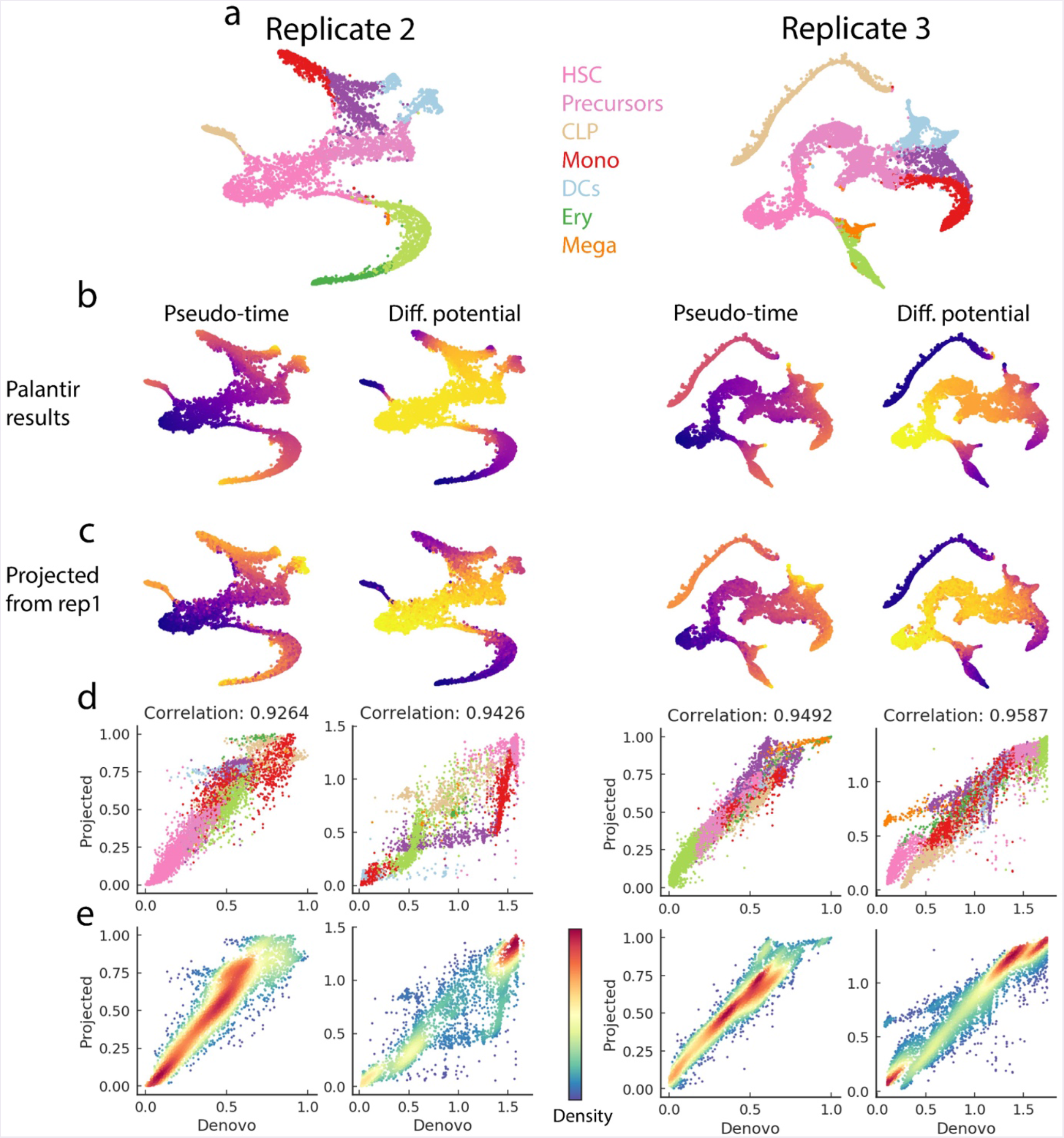
Reproducibility of Palantir pseudo-time and differentiation potential across replicates. (a) tSNE plots highlighting the different cell populations in replicate 2 (left) and replicate 3 (right). Cells are colored by Phenograph clusters and colors were chosen to maintain consistency with replicate 1 (Fig. 2b). (b) *De-novo* Palantir results for replicates 2 and 3 generated using one of the HSCs as the start cell. The reproducibility of Palantir results was further tested by projecting Palantir results of one replicate to a second replicate and comparing the projected results with those generated *de-novo* using the second replicate. (c) Replicate 1 Palantir results projected onto the replicates 2 and 3. The projections were determined mutually nearest neighbors between replicate 1 and replicate 2 (or 3), see methods. The pseudo-time and differentiation potential of cells in replicate 2 (or 3) were computed as weighted average of the pseudo-time and differentiation potential respectively of replicate 1 mutually nearest neighbors. (d) Plots showing the correlation between de-novo and projected Palantir results. The cells are colored by clusters as in (a). (e) Same as d, with cells colored by density.

**Supp. Fig. 8:**
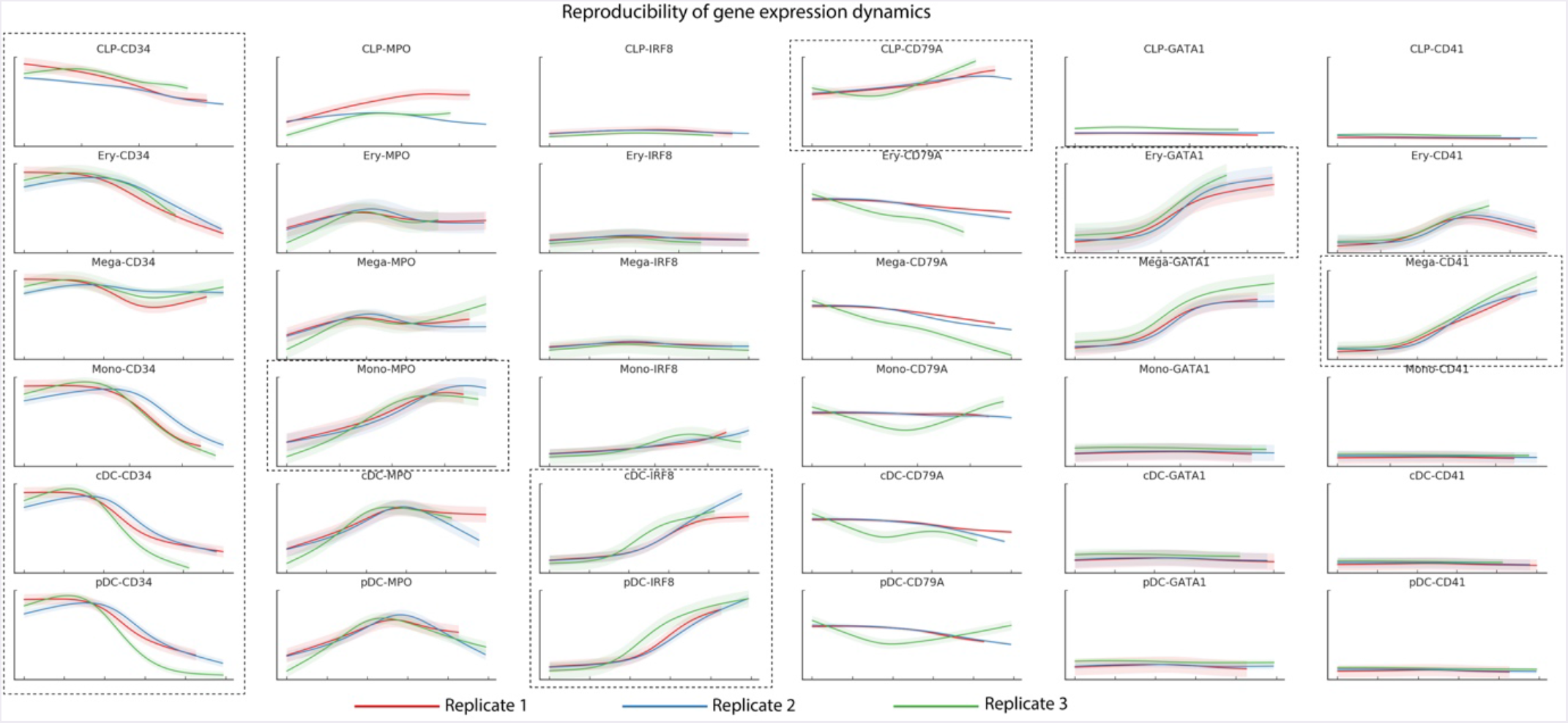
Reproducibility of gene expression dynamics across replicates. Plots showing reproducibility of expression trends of key lineage marker genes across the three replicates. The relevant lineages for each gene are bounded by dotted rectangles.

**Supp. Fig. 9:**
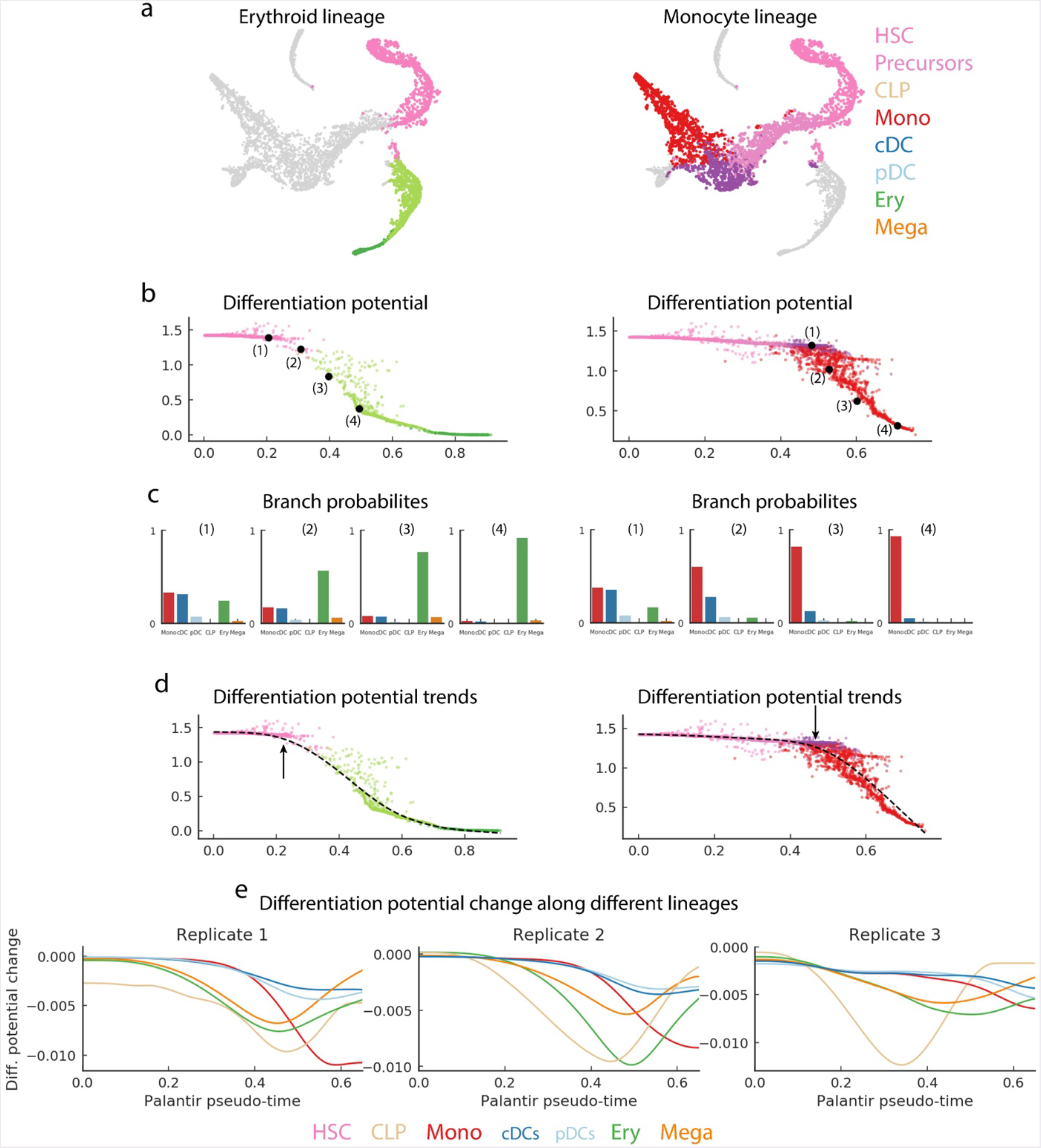
Illustration of Differentiation potential. (a) tSNE plots highlight the cells of the erythroid lineage (left) and monocyte lineage (right). Cells are colored by Phenograph clusters as in Fig. 2b. (b) Plots of DP (y-axis) along pseudo-time (x-axis). Each dot is a cell, color coded by the cluster. Representative cells are highlighted and numbered for each lineage. (c) Histogram representation of the branch probabilities of the cells highlighted in (b), bars are colored coded by clusters in Fig. 2b. As cells commit towards a particular lineage, they also lose the ability to differentiate to other lineages. This is reflected in the gradual increase in probability of reaching the corresponding terminal state accompanied by a decrease in probability of reaching all other lineages. (d) Same as (b), with trend plot of the DP along the corresponding lineages shown in black. The position along the ordering with the first substantial drop in DP corresponds to lineage specification and this downward trend continues until the cells are committed to the lineage and have completely lost the ability to differentiate to any other lineages. (e) Plots showing the DP along pseudo-time for all the lineages, for the three replicates. The positions of significant DP changes are staggered along the pseudo-time indicating a hierarchical commitment of HSCs to different lineages. Cells first commit towards CLP (beige), followed by erythroid and megakaryocytic lineages (green, orange) and finally the myeloid lineages: monocyte (red) and DCs (blues).

**Supp. Fig. 10:**
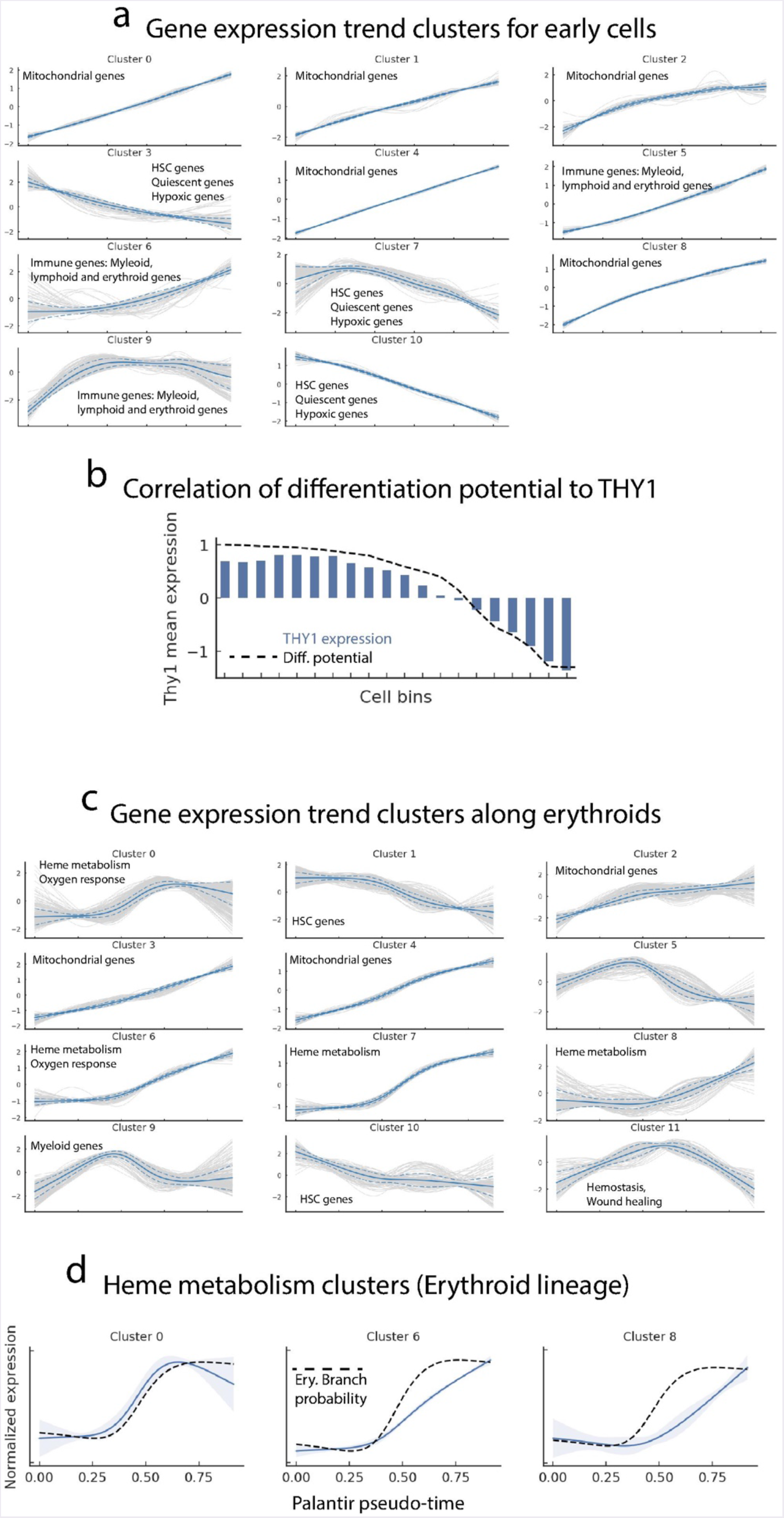
Clustering of gene expression trends in early and erythroid cells. (a) Plots showing cluster results of expression trends of genes that are significantly high or low in early hematopoietic clusters (0 and 1 in Supp. Fig. 4a). Each grey line represents the expression trend of a particular gene. Solid blue line represents the mean expression trend of the cluster and dotted blues lines represent the standard deviation. Each panel is labeled with enriched gene ontology terms. (b) Similar to Fig. 3b with the bar plot representing the mean expression of Thy1 in the bins. (c) Same as a - for genes significantly high or low in the clusters that correspond to the erythroid lineage (clusters 0, 1, 2 and 8 in Supp. Fig. 4a). (d) Average gene expression trends of heme metabolism clusters (0, 6 and 8 in b). Erythroid branch probability shown in black. Cluster 0 genes correlates the most with erythroid branch probability and are enriched with key erythroid TFs. Cluster 6 genes include KLF3. Cluster 8 contains genes such as HBB, that confer functional identity to erythroid cells.

**Supp. Fig. 11:**
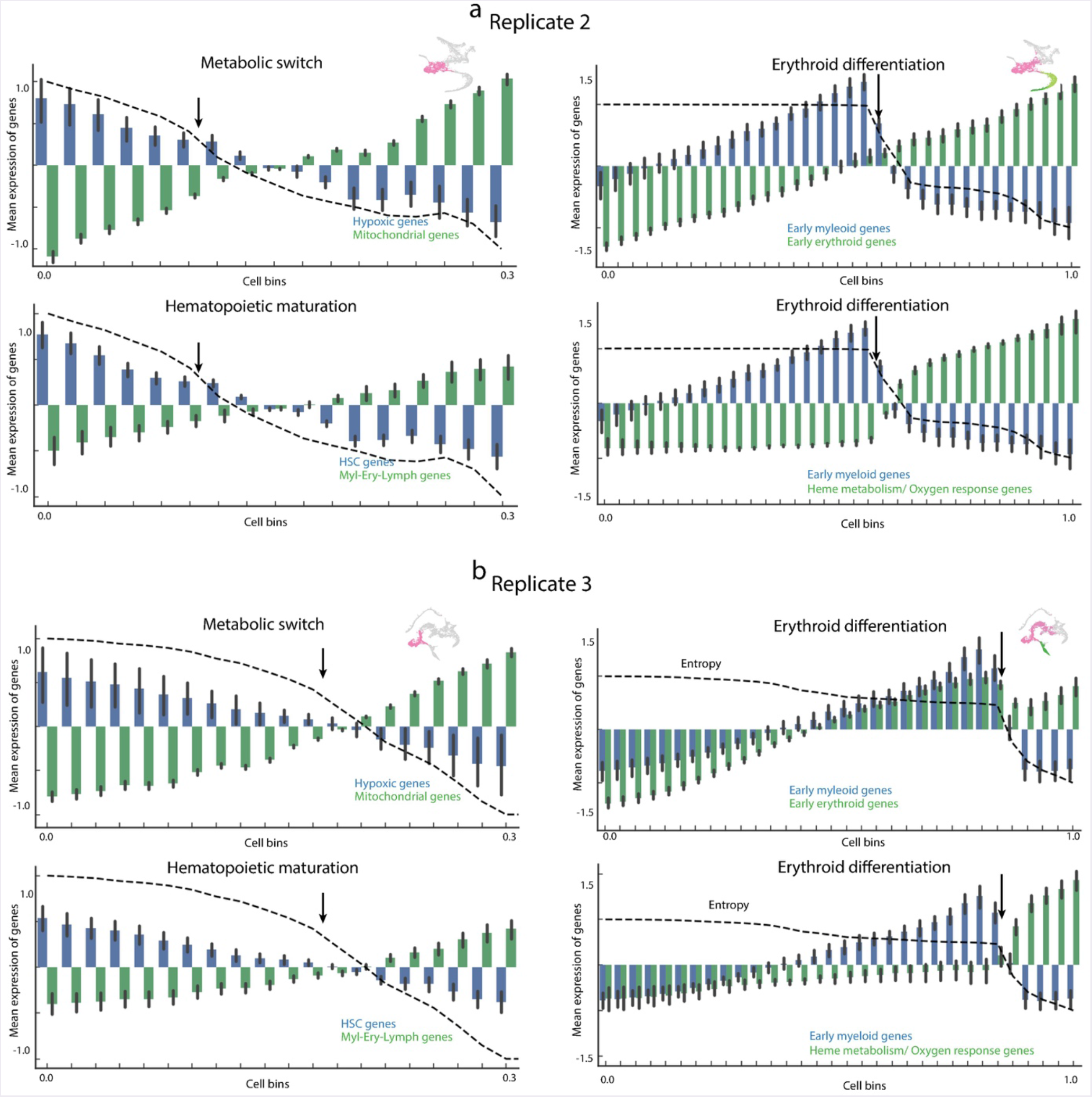
Palantir differentiation potential identifies landmarks of hematopoietic differentiation (Replication of results in Fig. 3) Plots showing the reproducibility of results in Fig. 3 in replicates 2 and 3. Genes identified using replicate 1 were used for this analysis.

**Supp. Fig. 12:**
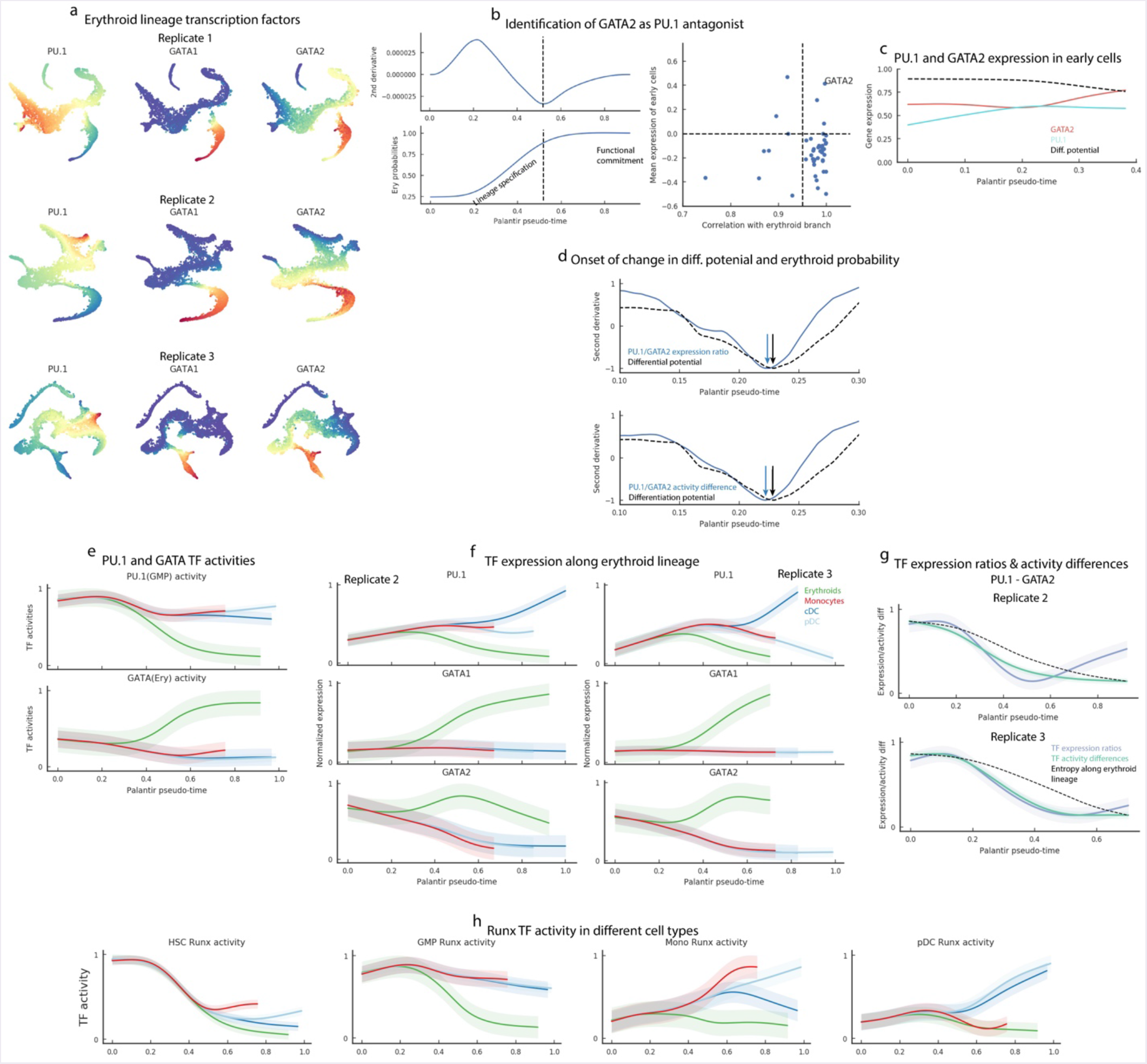
Identification of GATA2 as an agonist of PU.1 in driving erythroid specification. (a) tSNE plots showing the expression of PU.1, GATA1 and GATA2 in the three replicates. (b) Top left: Second derivative of erythroid branch probability trend. Dotted black represents the inflection point, i.e., the point of maximum change. Bottom left: Erythroid branch probability trend along Palantir pseudo-time. Right panel: Scatter-plot of TF expression trends with erythroid branch trends during the lineage specification phase (x-axis) and average TF expression in the early cells (y-axis). Each dot represents a TF. GATA2 (labeled) is a clear outlier. (c) Gene expression trends of PU.1 and GATA2 in early cells. Black line represents differentiation potential. (d) Top panel: Plot showing the second derivatives and the inflection points of the PU.1/GATA2 expression ratio (in blue) and the differentiation potential (in black). The change in expression ratio (blue arrow) precedes the change in differentiation potential (black arrow) indicating that PU.1/GATA2 ratio is predictive of lineage commitment. Bottom panel: Same as the top panel, but showing the TF activity differences between PU.1 and GATA2. (e) PU.1 and GATA1 activity trends along erythroid and myeloid lineages. PU.1 activities were determined using bulk GMP cells and GATA activities using bulk erythroid cells. PU.1 activity, is similar to its expression, showing an increasing trend in the myeloid lineages and GATA activity shows an upregulation specifically in the erythroid lineage. (f) Same as Fig. 4a, for replicates 2 and 3 (g) Same as Fig. 4c, for replicates 2 and 3 (h) Plots showing the Runx activity trends determined using Runx targets in different sorted populations. The activity is high specifically in the cell type from which the targets were derived highlighting the cell - type specificity of TF targets and the inferred TF activity.

**Supp. Fig. 13:**
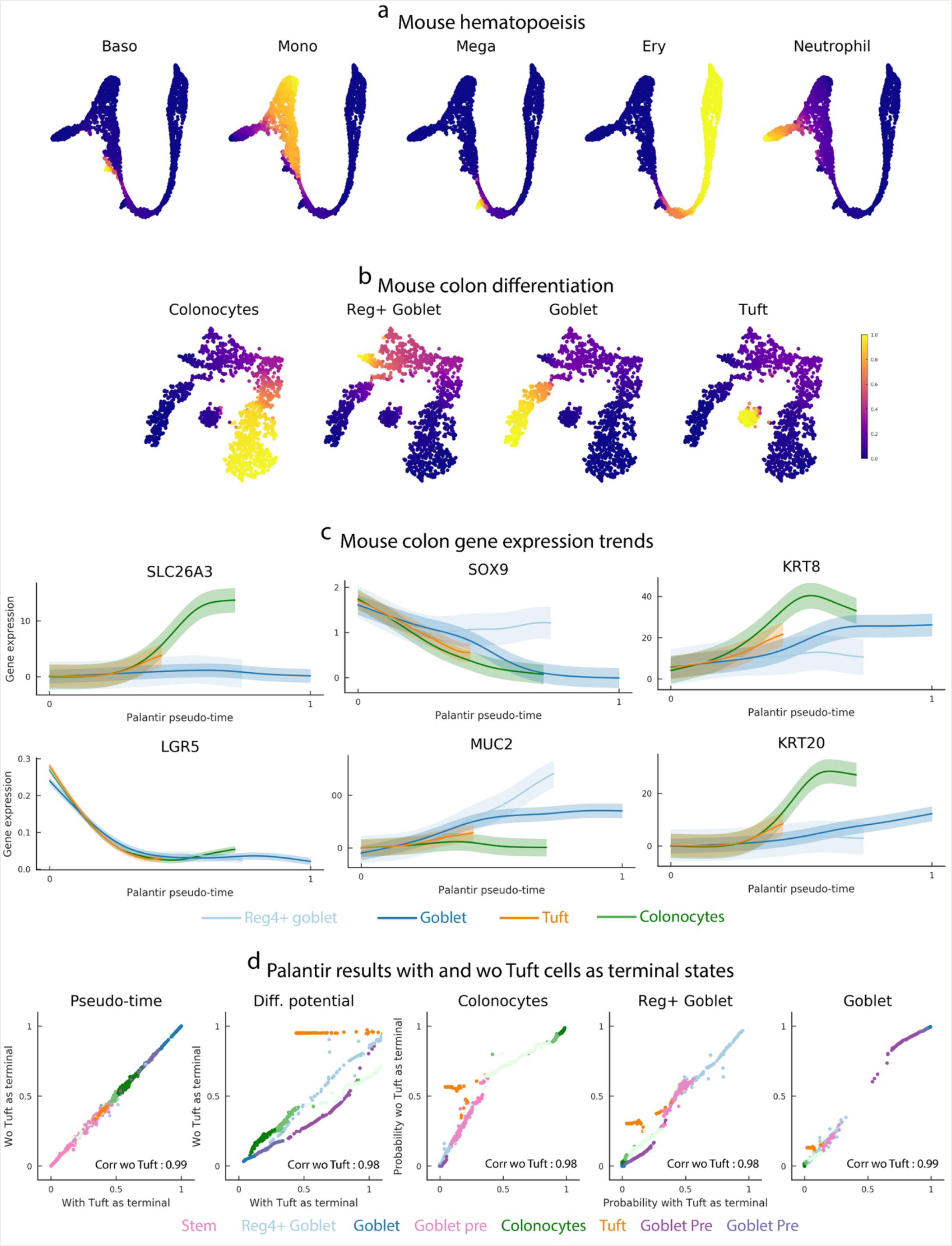
Palantir generalizes to mouse hematopoiesis and colon differentiation. (a-b) tSNE plots, with cells colored by Palantir branch probabilities for mouse hematopoiesis ^6^ and colon differentiation datasets ^48^ respectively. (c) Gene expression trends for the mouse colon data: Palantir results recapitulate the known behavior of key genes along different lineages. (d) Plots showing the correlation between Palantir results when Tuft cells (orange) were included (x-axis) or excluded as a terminal state (y-axis). Cells are color coded based on clusters in Fig. 5a.

**Supp. Fig. 14:**
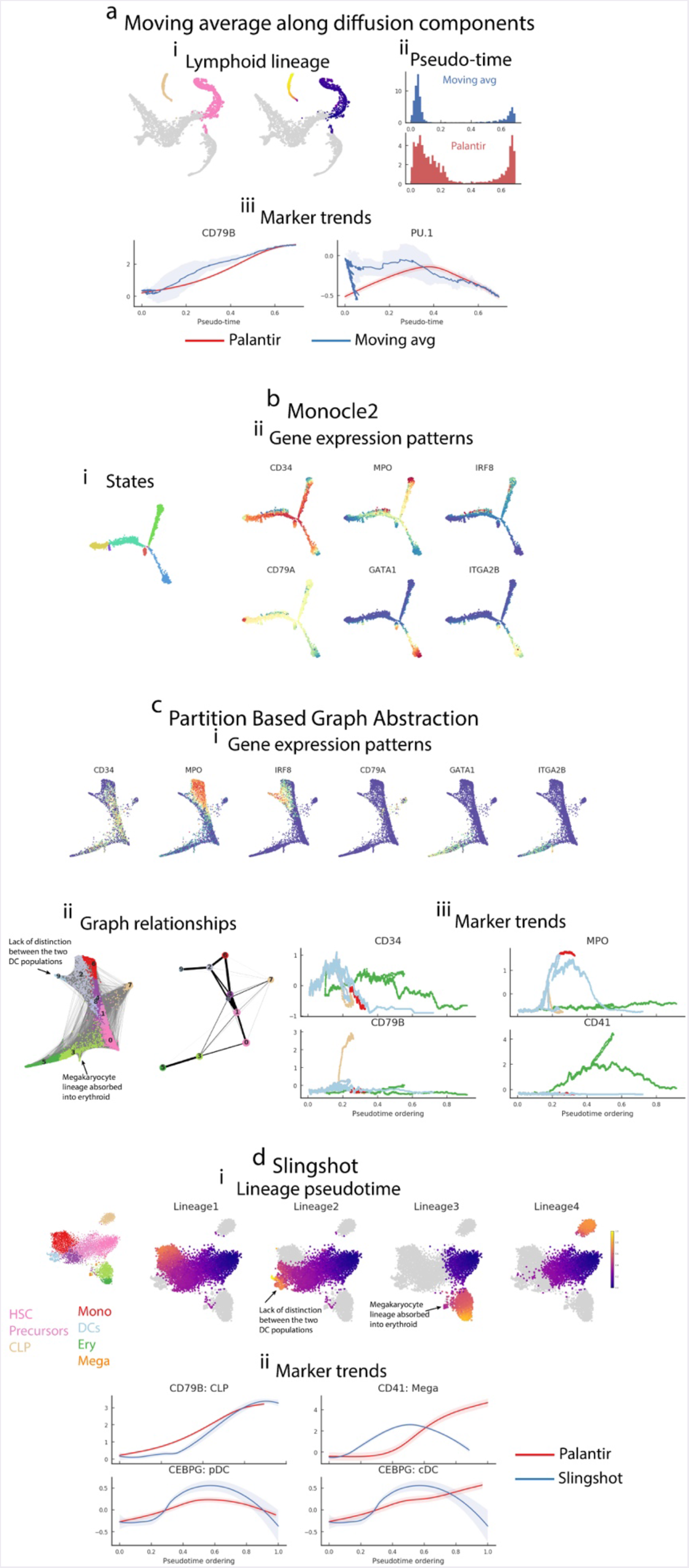
Performance of alternative pseudo-time methods on human hematopoiesis data. (a) Moving average along individual diffusion components: (i) (left) tSNE plots highlight the cells along lymphoid lineage and the pseudo-time of these cells as determined by projection along the 2^nd^ diffusion component (Supp. Fig. 2b). (ii) Histograms showing the density of distribution of cells along diffusion component (blue) or Palantir pseudotime (red). (iii) Gene expression trends determined by sliding windows (blue) and Palantir (red). (b) Monocle2 generated representation of the human hematopoiesis data, each dot is a cell (i) Cells colored by one of 6 Monocle 2 identified states. (ii) Cells colored by gene expression of key lineage genes. (c) Partition based Graph Abstraction (PAGA) applied to human hematopoiesis data (i) PAGA representation of the data highlighting key marker genes. (ii) Left: Connectivity among the hematopoietic cells as inferred by PAGA. Each dot is a cell color coded to represent lineages in Fig. 2b. Grey lines represent edges between two cells. Right: PAGA abstract clusters color coded to resemble Phenograph clusters in Fig. 2b. Thickness of the connection between clusters represents the strength of connectivity. PAGA lacks distinction between the two DC lineages and embeds megakaryocyte lineage to be part of erythroid lineage. (iii) Gene expression trends of key lineage markers as determined by PAGA. The trends are computed using a sliding window, making the estimates highly sensitive to noise in the data. (d) Slingshot applied to human hematopoiesis data, results are presented on tSNE maps generated using principal components of the data. (i) Cells colored by Slingshot pseudo-time for the four different lineages Slingshot identified. Slingshot results lacks distinction between the two DC lineages and embeds megakaryocyte lineage to be part of erythroid lineage. (ii) Gene expression trends determined by Slingshot (blue) and Palantir (red) with Slingshot trends showing (1) unexpected marginal downregulation of CD79B at the beginning of CLP lineage. (2) unexpectedly high upregulation of CD41 along the erythroid lineage since the megakaryocytes are included as part of the erythroid lineage and (3) lack of distinction between CEBPD dynamics in the two DC lineages since they were embedded as part of the same lineage.

**Supp. Fig. 15:**
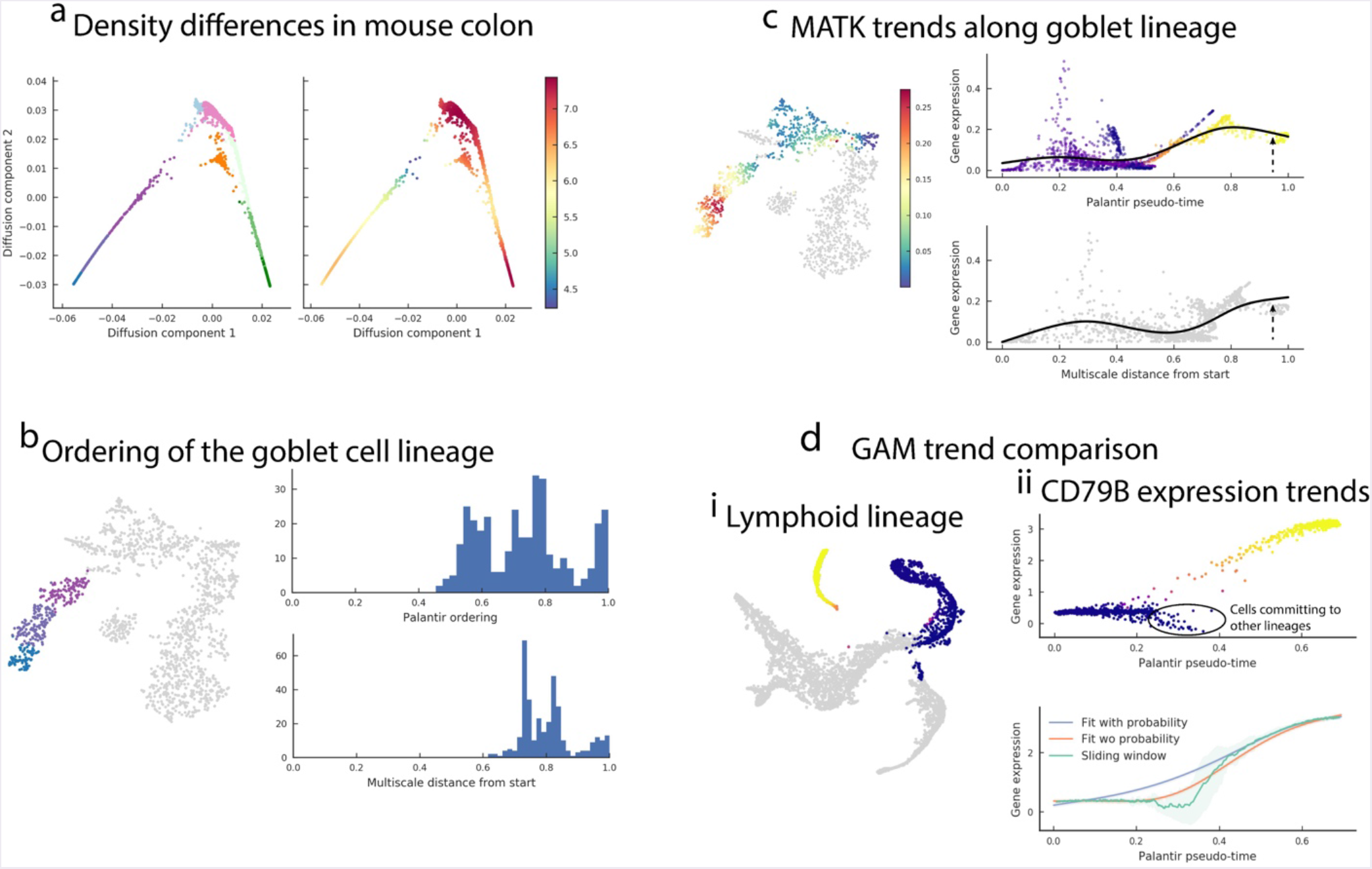
Palantir features key to identifying accurate identification of pseudotime ordering and branch probabilities. (a) Projection of the mouse colon data (Fig. 5) along the first two diffusion components. Each dot it a cell and is color coded by clusters in Fig. 5a (left panel) and density (right panel). (b) (i) tSNE map highlighting cells of the goblet lineage. Cells are color coded by clusters in Fig. 5a. (ii) Histograms showing the distribution of goblet cells along the pseudo-time derived using shortest path distances (top panel) and multi scale distances (bottom panel). Distribution of cells using multiscale distance leads to loss in resolution with increased local concentration of goblet cells at the end of the pseudo-time. (c) (i) tSNE plot showing the expression of MATK along the cells that contribute to the goblet cell lineage (left panel). (ii) Plots showing MATK expression along pseudo-time where each dot is a cell. Top: Cells colored by goblet cell probability. Black line represents the MATK trend obtained by GAM fit using goblet cell probabilities as weights for cells. Bottom: MATK trend with GAMs fitted on cells ordered by pseudo-time generated using multi scale distances. The loss of resolution in pseudo-time results in even GAMs miss trends such as MATK downregulation towards the end of goblet lineage. (d) (i) tSNE plot highlighting the cells along the lymphoid lineage (clusters 0 and 5 in Supp. Fig, 4a) colored by lymphoid branch probability from the human hematopoiesis dataset. (ii) Top: Plot showing expression of CD79B along Palantir pseudo-time. Each dot is a cell and is colored by lymphoid branch probability. Cells that are committing towards other lineages are highlighted. Bottom: Gene expression trends computed using sliding windows (green), GAMs fit without using Palantir probabilities (orange) and GAMs fit using Palantir probabilities (blue). Sliding window and GAMs fit without using branch probabilities are heavily influenced by cells committing to other lineages leading to incorrect trend estimates.

**Supp. Fig. 16:**
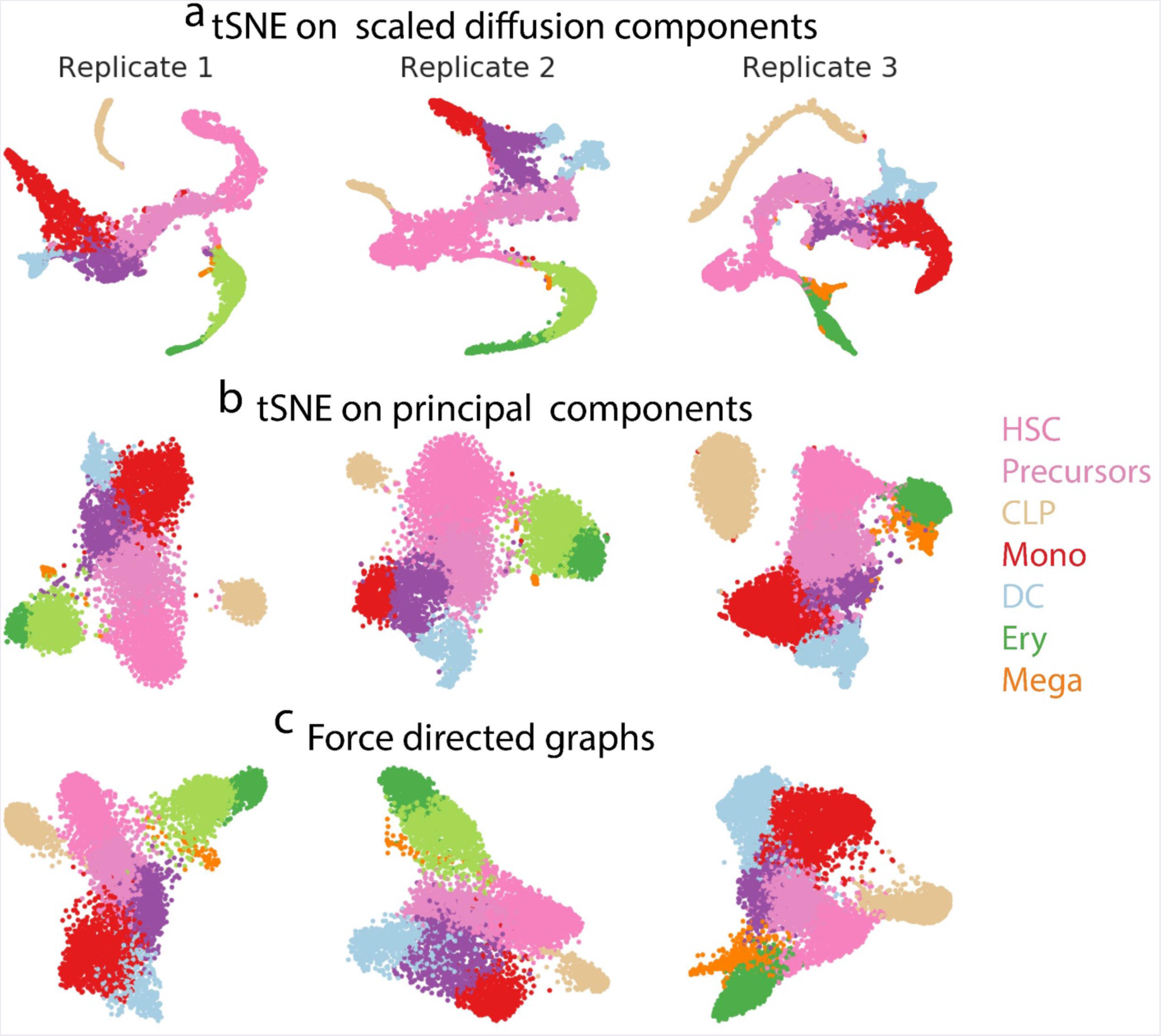
Comparison of data visualizations. tSNE maps generated using (a) scaled diffusion components and (b) principal components. (c) Force directed graphs for the same cells. Cells are color coded by clusters in Fig. 2b.

**Supp. Fig. 17:**
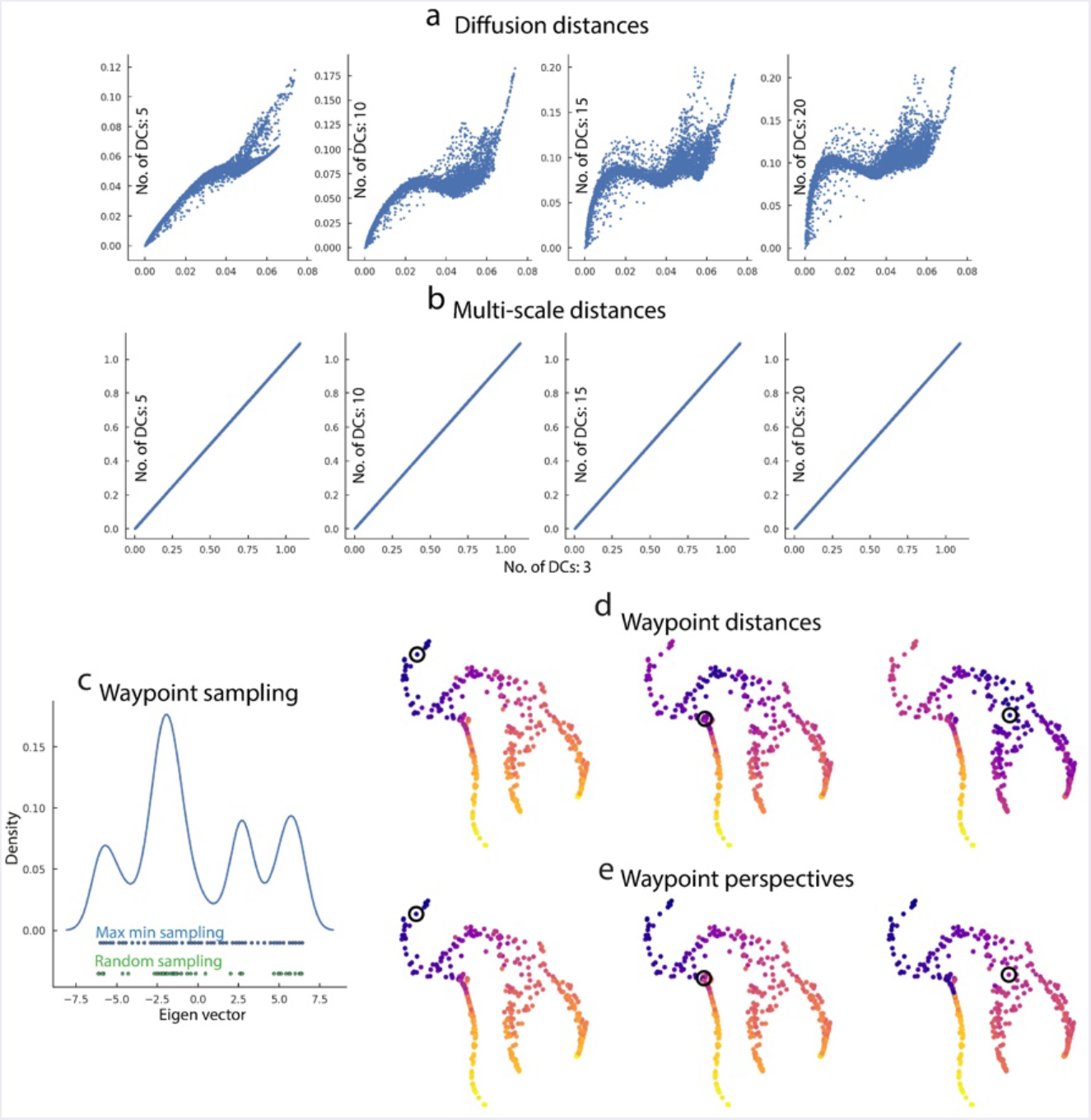
Multi-scale distances, waypoint sampling and perspectives. (a) Plots comparing the diffusion distances from an early HSC cell when different number of components are used. (b) Same as (a), with distances computed using multi-scale distance. (c) Plot showing the variable density of cells along a particular diffusion component. Random sampling of waypoints samples cells from high density regions while ignoring of low density and high variability (green dots). Max-min sampling however samples cells along the entire spectrum of the diffusion component and generates a more representative sample of cells (blue dots). (d) Shortest path distances from a subset of waypoints. (d-e) Cells from subsampled dataset (Fig. 1b) colored by shortest path distances and perspectives from the highlighted waypoints.

**Supp. Fig. 18:**
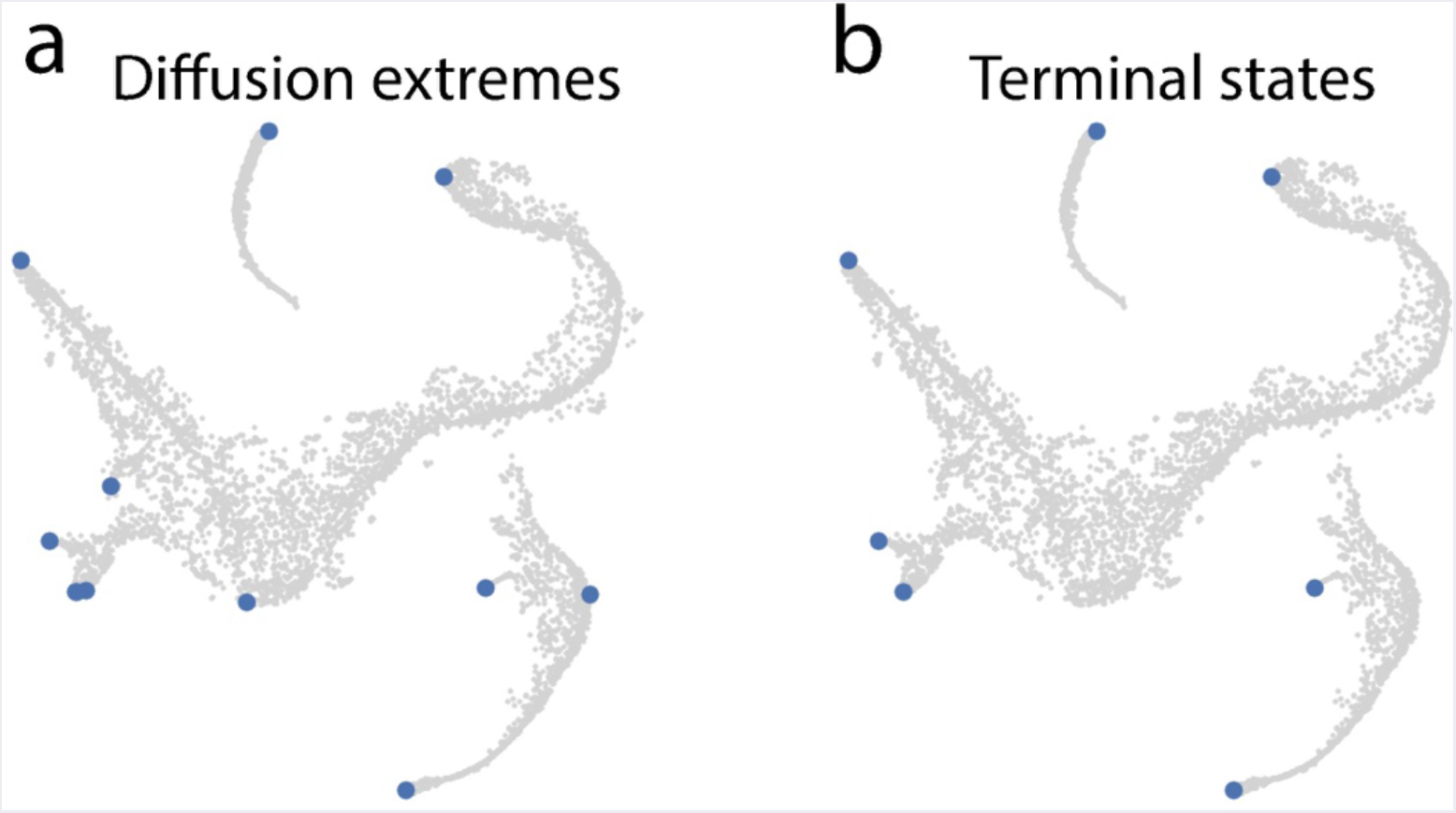
Identification of terminal states. (a-b) Diffusion map boundaries and the identified terminal states for replicate 1 of the human hematopoiesis data.

